# Causality-oriented regulatory inference and rational transcriptomic reprogramming design with CASCADE

**DOI:** 10.1101/2025.11.20.689377

**Authors:** Zhi-Jie Cao, Tianzi Guo, Cheng Li, Fan Zhang, Ping Lu, Chan Gu, Fan Guo, Ge Gao

## Abstract

Systematically elucidating the causal structure of gene expression regulation is the foundation for identifying key regulators of cellular function and designing rational interventions thereof. While advances in high-content perturbation screens enables experimental probing of causal regulatory effects, the massive genomic scale, measurement noise and confounding factors pose significant challenges for existing causal discovery and inference algorithms. Here, we present CASCADE, an end-to-end framework that integrates *de novo* causal discovery with prior regulatory knowledge for genome-wide causal discovery and inference. In addition to its superior performance on causal regulatory structure discovery and counterfactual inference for unseen perturbation effects, CASCADE enables, for the first time, causality-oriented rational intervention design for transcriptomic programming towards various targeted cells within a unified framework.

## Introduction

Gene expression regulation is the basis of how genetically identical cells differentiate into diverse cell types with distinct morphologies and functions^1,2^. A genome-scale causal understanding of the gene regulatory network (GRN) is essential for precise *in silico* prediction of global cellular intervention response and rational cell programming design, and the cornerstone of virtual cells^3–5^,

While high-content single-cell perturbation screens such as Perturb-seq allow for experimentally probing causal regulatory effects^5^, the impractical nature of exhausting the combinatorial perturbation space calls for causality-oriented computational methods capable of generalizing to unmeasured perturbations through discovering the causal GRNs underlying the observed outcomes^6^. Meanwhile, the scale of thousands of genes in eukaryotic cells effectively overwhelms canonical causal structure discovery methods^7–12^, and the extensive noise and confounding factors in current single-cell data further pose serious challenges for precise identification of genuine causal relations^13^, much less utilizing the inferred causal relations for effective perturbation effect prediction and design.

Combining *de novo* causal discovery with prior regulatory knowledge, we propose **<u>CASCADE</u>** (**<u>C</u>**ausality-**<u>A</u>**ware **<u>S</u>**ingle-**<u>C</u>**ell **<u>A</u>**daptive **<u>D</u>**iscover/**<u>D</u>**eduction/**<u>D</u>**esign **<u>E</u>**ngine), the first end-to-end regulatory causality inference and design model at genome scale. Based on the inferred network, the model achieves interpretable counterfactual inference of unseen single-gene and multi-gene perturbation responses through iterative effect propagation, and finally, empowers automatic intervention design to achieve desired effects of targeted cell programming (Fig. 1A). CASCADE is publicly available at https://github.com/gao-lab/CASCADE.

**Fig. 1.**
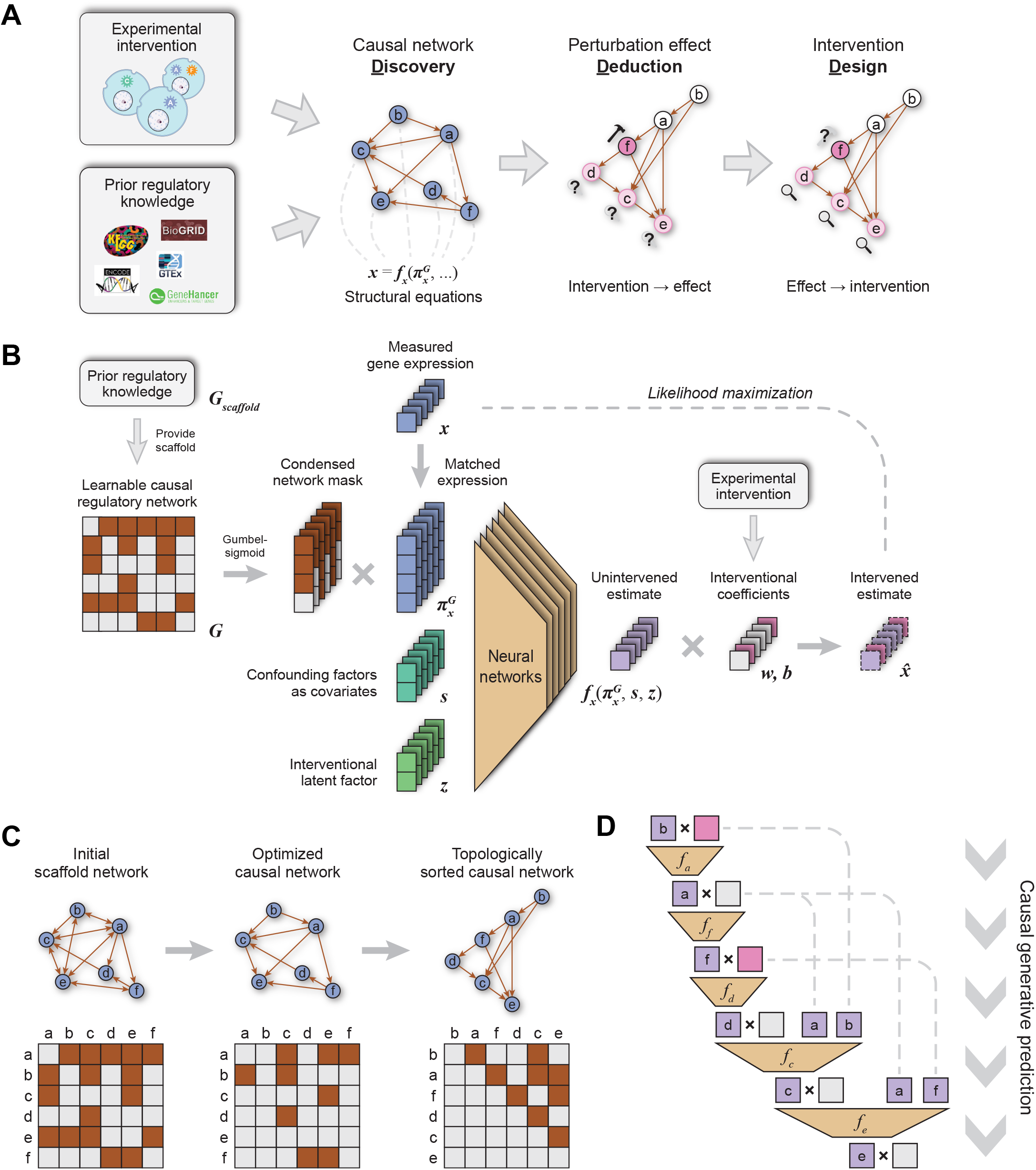
Schematic of the CASCADE model and causal inference procedures. (**A**) General capability of the CASCADE model. (**B**) Architecture of the CASCADE model. The model explicitly optimizes the adjacency matrix of GRN **G** starting with a scaffold graph **G**_scaffold_ built from heterogeneous prior regulatory knowledge. Each gene is assigned an MLP neural network with GRN-masked parent genes as input, which outputs an estimate of its unintervened expression. This estimate then undergoes an interventional transformation to produce the final estimate 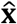. Confounding factors such as batch effect and sequencing depth are modeled as covariates **s**. A latent factor **z** inferred from GO-based functional prior knowledge is also included to fit residual effects. The entire model is trained using self-reconstruction likelihood maximization. (**C**) During causal discovery, edges in the scaffold graph are pruned to orient undirected edges and break cycles, producing a causal DAG that best explains the data, based on which all genes can be topologically sorted. (**D**) The gene-wise MLPs can then be unrolled based on the topological order into a sequential causal generative model. Counterfactual perturbation effects are predicted by manipulating the perturbation label for each gene and allowing the resulting effect to propagate throughout the model.

## Results

### Causality-oriented gene regulatory modeling

By modeling the genome-scale causal GRN as a learnable directed acyclic graph (DAG), CASCADE quantifies causal regulatory interactions as non-linear structural equations (Fig. 1A)^14^. Specifically, each gene is assigned a neural network structural equation (Fig. 1B, center) that predicts its unintervened expression from upstream regulators (determined by GRN adjacency, Fig. 1B, left), along with multiple confounding factors such as batch and sequencing depth (Methods). The unintervened estimates of intervened genes are then transformed to fit the observed expression based on a set of learnable interventional coefficients (Fig. 1B, right). Both the causal GRN and the structural equations are optimized using differential causal discovery (DCD) to minimize reconstruction error. Inspired by previous work^15^, CASCADE supports a scaffold graph compiled from context-agnostic, coarse prior regulatory knowledge to constrain the GRN search space in an evidence-guided manner (Fig. 1B, top left). Furthermore, we encapsulate regular DCD under a Bayesian framework based on Stein variational gradient descent (SVGD), allowing for the estimation of causal uncertainty under limited data regimes typical of practical biological experiments (Methods).

Using the inferred causal GRN, CASCADE supports two types of downstream inference. First, it performs counterfactual deduction of unseen perturbation effects by iteratively propagating perturbation effects following the topological order of the causal graph (Fig. 1B, C). Notably, this deduction process remains end-to-end differentiable, allowing it to be inverted into intervention design by treating gene intervention as an optimizable parameter trained to minimize deviation between the counterfactual outcome and desired target transcriptomes (Methods).

### CASCADE is accurate for genome-scale casual discovery

We first adopted a simulative approach where causal discovery methods were applied to data simulated from known DAGs, with normalized marginal variances to prevent “varsortability”^16^ (Methods). Among all methods tested, CASCADE demonstrated the highest accuracy across a range of gene numbers, intervened gene fractions, as well as different evaluation metrics (Fig. 2A, Fig. S2A, Table S2), with a more pronounced advantage at larger scales (Fig. 2B, Fig. S1). Consistent with their designs that explicitly model interventions, GIES, DCDI and CASCADE showed the most pronounced improvement with increasing intervention data incorporated (Fig. 2A).

**Fig. 2.**
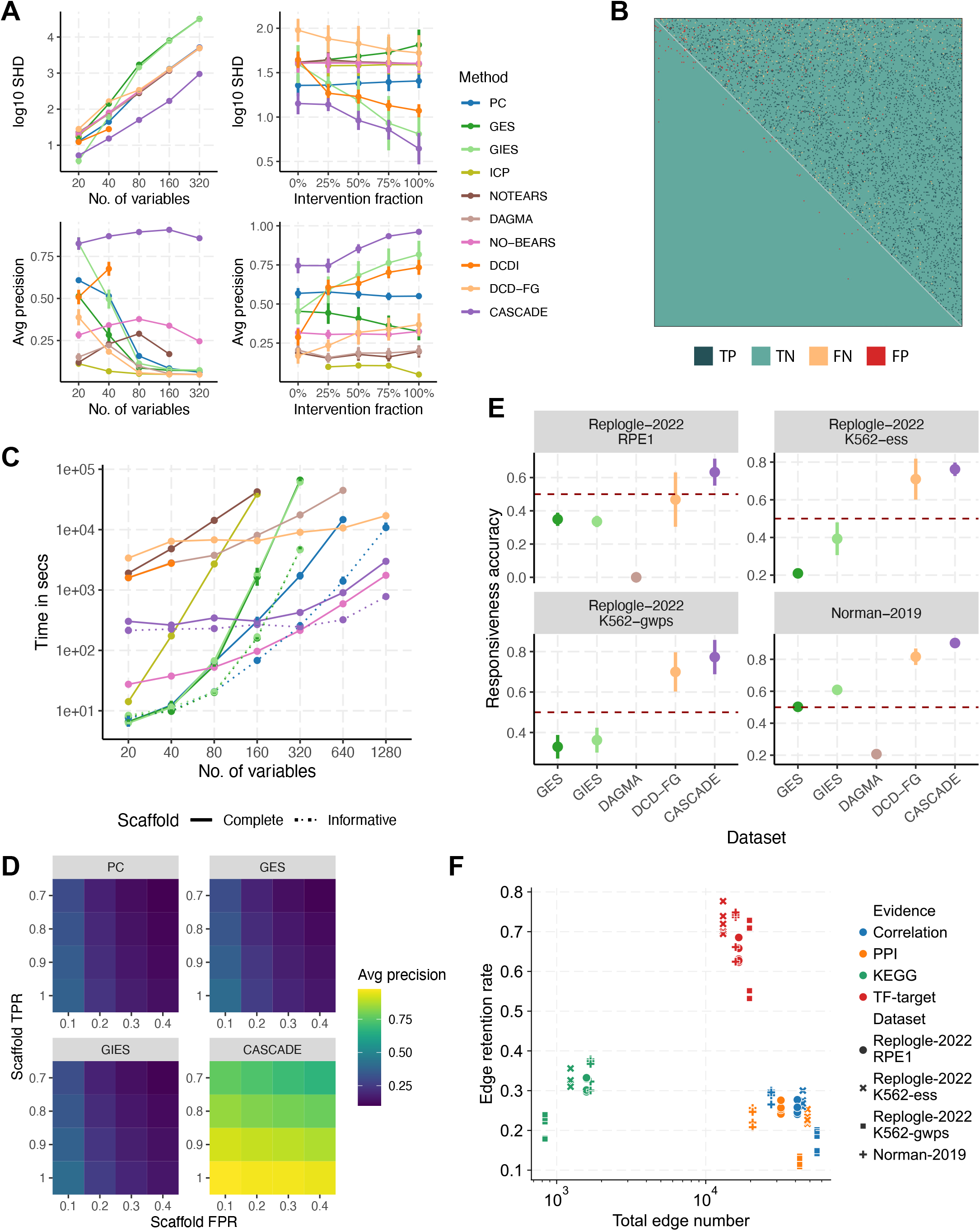
Systematic benchmarking of causal discovery accuracy. (**A**) Ground truth accuracy of the causal networks inferred by different methods from simulated data, as quantified by log_10_ structural Hamming distance (SHD) and average precision, under varying numbers of variables and fractions of intervened variables. The intervention fraction plots only summarize results from 20 and 40 variables due to DCDI failing beyond 80. N = 4 with different true DAGs. Error bars indicate mean ± s.d. (**B**) An example CASCADE-inferred adjacency matrix at 320-variable scale. TP, true positive; TN, true negative; FN, false negative; FP, false positive. (**C**) Running time of different causal discovery methods with up to 1,280 variables and 50% intervened fraction. The informative scaffold was obtained by corrupting the true DAG with 10% false positive edges. N = 4 with different true DAGs. (**D**) Accuracy of different methods when using scaffold graphs of varying TPRs and FPRs, as quantified by average precision within scaffold edges. (**E**) Performance on different Perturb-seq datasets at a 200-gene scale, as quantified by responsiveness accuracy. N = 5 with different train/test splits. (**F**) The fraction of scaffold edges in each evidence type retained after model training.

Notably, biological networks easily contain thousands of genes, substantially larger than even the largest scales tested above. While many canonical methods such as GES and GIES performed well at small numbers of genes, their performance declined significantly as the simulation scale increased to hundreds of genes (Fig. 2A, Fig. S1). Of note, while DCDI exhibited a promising performance trend, its inferior scalability led to failed runs within a 24-hour time limit for just 80 genes. Only DCD-FG, NO-BEARS and CASCADE managed to process over one thousand genes, and CASCADE was the most accurate with a reasonable running speed (Fig. 2C and Fig. S2B, solid lines, Table S3, 4).

Causal discovery on real single-cell perturbation data can be more challenging than the simulative setting due to confounding effects, and more importantly, regulative mechanisms such as post translational modifications not directly assessed in the transcriptome assay. Thus, an interventional latent variable encoded from functional annotation of intervened genes was incorporated into CASCADE to account for effects not explained by the transcriptomics network and mitigate spurious causal edges (Fig. 1B, bottom center, Methods). Given the fact that the ground truth biological GRN is largely unknown, we sought to assess the real-world causal discovery performance based on the asymmetry of perturbation effects: for correctly inferred regulatory edges, perturbing the parent gene should produce greater effect on the child gene, rather than the other way around. Thus, a new metric, “responsiveness accuracy”, is introduced to quantify how well the inferred GRNs align with this assumption (Methods).

We used 4 Perturb-seq datasets covering both CRISPRi and CRISPRa, with single- and double-gene perturbations^17,18^ (Table S1). Among all methods that successfully completed at a 200-gene scale (Methods), CASCADE achieved the highest responsiveness accuracy across datasets (Fig. 2E, using the complete scaffold for fairness, Table S5), extending its advantage in simulative settings to real Perturb-seq data. The accuracy of CASCADE improves with multiple SVGD particles (Fig. S4A), indicating that modeling graph uncertainty is important in the complex and noisy gene regulatory system. CASCADE was otherwise robust across a range of hyperparameter settings (Fig. S4A, Table S6).

While the scaffold graph was disabled in all the above benchmarks entirely (for fair comparison with methods that cannot utilize scaffolds), we sought to assess the utility of scaffold graphs further, first with simulated data. To reflect the reality that prior regulatory knowledge is often inaccurate, informative yet inaccurate scaffold graphs were constructed by corrupting the ground truth DAGs with 10% false positive edges. Consistently, we found incorporating scaffold graphs effectively improved performance (Fig. 2C, and Fig. S2B, dotted lines, Table S3, 4), with CASCADE as the fastest and most accurate one. To assess the models’ robustness to various levels of prior knowledge inaccuracies, we tested them using scaffold graphs of varying true positive rates (TPRs) and false positive rates (FPRs). CASCADE demonstrated remarkable robustness to different numbers of false positive scaffold edges, particularly when TPR is high (Fig. 2D, Fig. S3, Table S7), while others exhibited much lower accuracy.

For real-world perturbation data, we constructed an informative but context-agnostic scaffold graph by combining heterogeneous types of evidence, including KEGG pathways^19^, TF-target relations from chromatin binding^20,21^, protein-protein interaction from BioGRID^22^, as well as highly correlated genes among GTEx tissues^23^ (Methods), which enables CASCADE to conduct causal discovery at the scale of 2,000 genes. We compared the scaffold edge retention rate of different evidence types after model training. Across all 4 datasets, TF-target edges exhibited the highest retention rate, followed by KEGG edges (Fig. 2F). Additionally, CASCADE predominantly oriented undirected scaffold edges in the TF-to-target direction (Fig. S4B), consistent with the understanding that TF regulation is the primary causal mechanism governing gene expression.

### CASCADE precisely models complex perturbation effects with interpretability

A causal GRN provides the reliable basis for CASCADE to predict perturbation effects which are never measured experimentally (i.e., counterfactual prediction, Methods). We first compared CASCADE with existing methods^24–26^ in predicting the effects of held-out single-gene perturbations. E.g., in the “Replogle-2022-K562-ess” dataset, CASCADE produced more accurate predictions for the top 20 differentially expressed genes (DEGs) upon *RPS9* knockdown compared to other methods (Fig. 3A). Systematic benchmarking involving 5 Perturb-seq datasets^17,18,27^ (Table S1) demonstrated that CASCADE consistently outperformed other methods in predicting held-out single-gene perturbation effects across different numbers of DEGs, quantified by both delta correlation and normalized mean square error (MSE) (Fig. 3C, Table S8, Methods), including large foundation models^28–30^ and a linear baseline which existing deep learning methods failed to surpass^31^. Ablation experiments also confirmed that the causal GRN contributed more than the latent factor (Fig. S5A, Table S9), highlighting the advantage of causal modeling over latent-based strategies in gene-wise perturbation generalization.

**Fig. 3.**
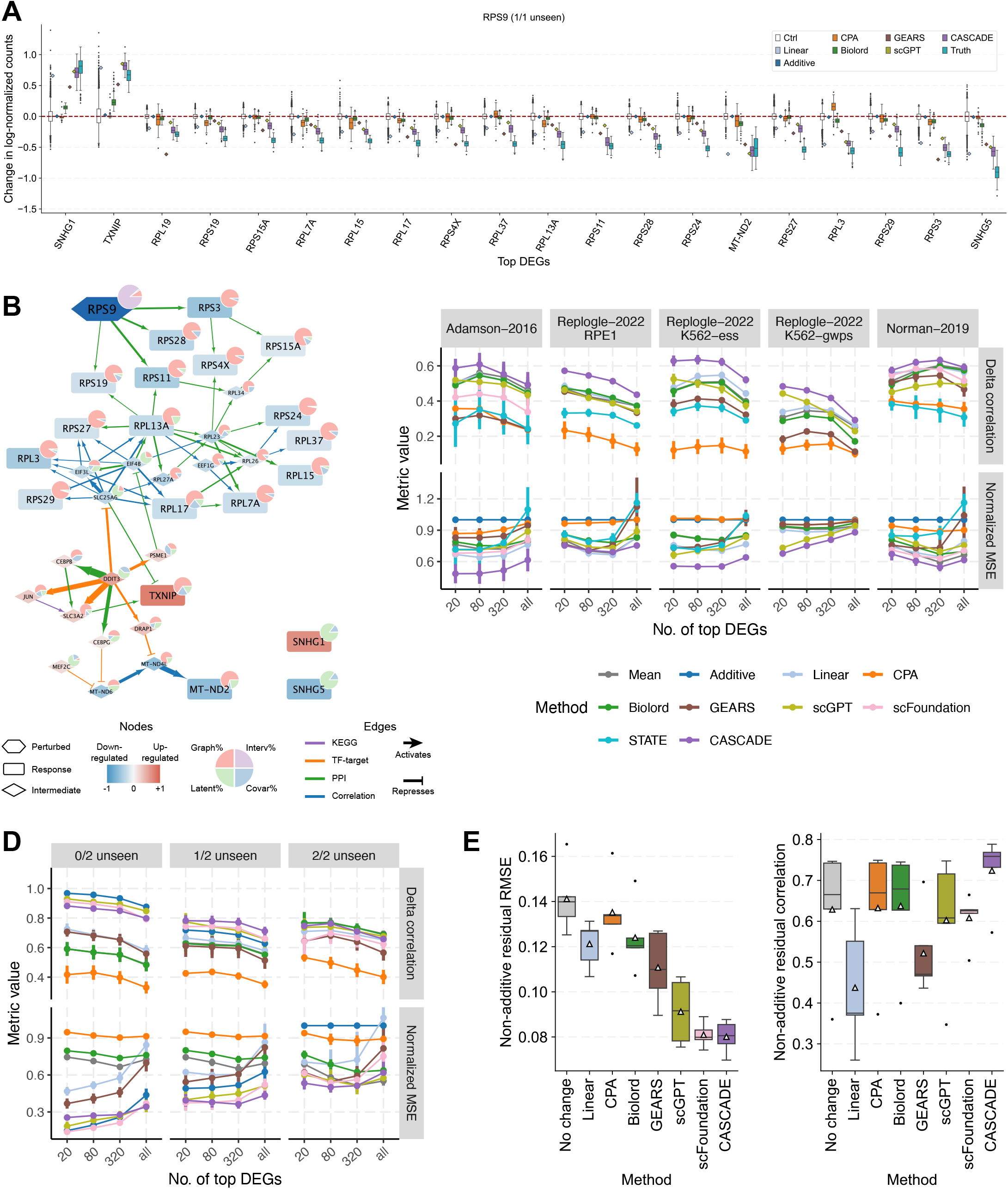
Counterfactual prediction of perturbation effects based on the causal model. Prediction comparison for an example perturbation in “Replogle-2022-K562-ess”, including predicted and true single cell expression distributions for the top 20 DEGs upon *RPS9* knockdown. Gene expressions were normalized by subtracting the median values in control cells. Box plots indicate medians (centerlines), first and third quartiles (bounds of boxes) and 1.5× interquartile range (whiskers). Core GRN explaining the counterfactual prediction in (**A**). Node colors indicate predicted expression changes. Pie charts indicate contribution proportions. Edge colors indicate scaffold evidence types. Edge widths indicate contribution amount. (**C**) Accuracy of single gene perturbation effect prediction for different methods on held-out perturbations across 5 Perturb-seq datasets, as quantified by delta correlation and normalized MSE among different numbers of DEGs. N = 5 with different train/test splits. Error bars indicate mean ± s.d. STATE failed on the “Replogle-2022-K562-gwps” dataset due to 24-hour training timeout. Of note, CPA^24^ and STATE^30^ were designed for cell context generalization, i.e., predicting perturbations seen during training in different cellular contexts than the test data. Our evaluation on predicting completely unseen perturbations is considerably more challenging, which may explain their underperformance. (**D**) Accuracy of double gene perturbation effect prediction on held-out perturbations in “Norman-2019”. N = 5 with different train/test splits. STATE could not be included because it lacks the ability to predict multi-gene perturbations. (**E**) Accuracy in predicting non-additive residual effects for 0/2 unseen perturbations, as quantified by Pearson correlation and RMSE. N = 5 with different train/test splits. Box plots indicate medians (centerlines), mean (triangles), first and third quartiles (bounds of boxes) and 1.5× interquartile range (whiskers).

Of note, benefitting from explicit modeling for the causal regulatory structure, CASCADE predictions are highly interpretable. We could attribute the predicted expression changes of each gene to covariates (experimental batch and total RNA counts), latent factor, direct CRIPSR perturbation, and regulation by upstream genes, respectively (Fig. 3B, Methods). E.g., among the 20 top DEGs caused by *RPS9* knockdown, only ribosomal genes (*RPL19, RPS19*, etc.) were attributed as its direct consequence, while the upregulation of *TXNIP*, and downregulation of mitochondrial respiratory chain genes like *MT-ND2* were linked to the activation of *DDIT3*, indicating a potential apoptotic response caused by ribosomal disruption (Fig. 3B).

For more complex double-gene perturbations, the methods could be tasked to predict the effect of unseen combinations where the constituent genes are both individually seen in training (0/2 unseen), one seen in training (1/2 unseen) or neither seen in training (2/2 unseen). CASCADE predictions for top DEGs could also be deconvoluted into direct and latent ones, with direct effects predicted more accurately (Fig. S6, Fig. S7), and functionally relevant genes showing more overlap in immediate downstream targets (Fig. S7). Benchmarks with different numbers of DEGs indicated that CASCADE, scGPT and scFoundation were close to an additive baseline in the 0/2 unseen category due to the dominance of additive effects in this dataset (Fig. 3D). However, CASCADE emerged as the most accurate, as quantified by Pearson correlation or root mean squared error (RMSE), when focusing on non-additive effects (Fig. 3E, Fig. S8), and also showed advantages in categories involving at least 1 unseen perturbation (Fig. 3D), with more contribution from the causal graph than the latent factor (Fig. S5B, Table S9).

The robustness of CASCADE was also evaluated under various hyperparameter settings. The hyperparameters that most significantly affect counterfactual accuracy are KL weight *β* and latent dimension *Z* (Fig. S9, 10, Table S10, Methods). Smaller *β* or larger *Z* increases the model’s latent capacity, which improved accuracy in tasks such as 0/2 unseen prediction but led to overfitting in the more challenging 2/2 unseen prediction (Fig. S10), illustrating the limitation of latent-based approaches. Our default setting (*β* = 0.1, *Z* = 16) limits latent capacity and encourages the model to rely more on the causal GRN. CASCADE remains robust to other hyperparameters.

### CASCADE enables rational intervention design for transcriptomic programming

Accurate prediction of complex perturbations implies an opportunity for designing optimal interventions for programming cells into desired states. While existing perturbation prediction methods can theoretically accomplish this by exhausting the effect of all potential perturbations^28^, CASCADE achieves this through a dedicated end-to-end differentiable optimization algorithm (Methods). Briefly, the identity of intervened genes and their interventional coefficients are treated as learnable model parameters, trained to minimize the deviation between the counterfactual outcome and a specific target transcriptome (Methods). By not requiring counterfactual prediction of every cell under every candidate intervention, the optimization-based approach is not only more efficient over canonical exhaustive search, but also enables a direct estimation of optimal intervention coefficients (rather than relying on fixed extrapolation from the training data).

We first assessed the design accuracy, in terms of identifying the true perturbed genes or gene combinations from their inflicted effects, using held-out perturbations in the 5 Perturb-seq datasets^17,18,27^ (Table S1). Inspired by the hit rate metric for reverse perturbation prediction^28^, we propose a new performance metric, area under hit rate curve (AUHRC), to assess design accuracy (Methods, Fig. S11A). Benefitting from its precise counterfactual prediction, CASCADE demonstrated the highest area under hit rate curve (AUHRC) in 4 out of 5 datasets under both exhaustive search and differentiable optimization modes (Fig. 4A, B, Table S11). Notably, optimization-based CASCADE designs were more accurate than exhaustive search due to its adaptive intervention coefficient estimation, which results in improved designs when the training extrapolations are markedly inaccurate (Fig. S11B).

**Fig. 4.**
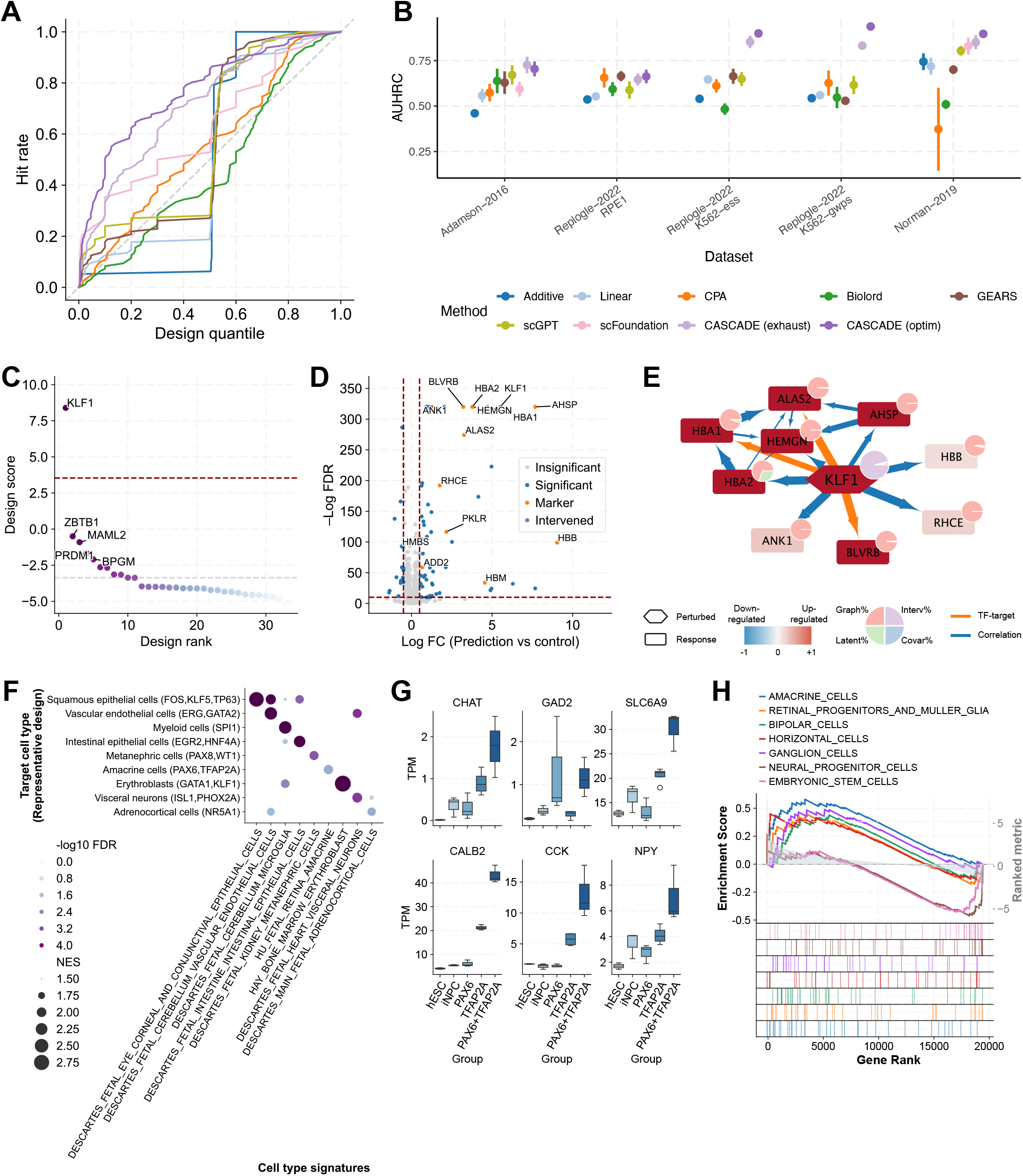
Rational intervention design guided by the causal model. (**A**) Hit rate curve (HRC) for different intervention design methods in identifying the perturbed genes/gene combinations given their inflicted effects. Design quantile indicates the quantile of the true perturbation among all possible perturbations ranked from highest to lowest scores. Hit rate indicates the fraction of true perturbations that would be covered at specific design quantiles. Larger area under HRC indicate more accurate designs (N = 125 from 5 datasets × 5 train/test splits × 5 designed perturbations). (**B**) Performance of intervention design across different datasets, as quantified by the area under hit rate curve (AUHRC). N = 5 with different train/test splits. Error bars indicate mean ± s.d. (**C**) Top ranked genes in CASCADE design for erythroid differentiation. Red horizontal line indicates the empirical design score cutoff covering the smallest MSE within 95% confidence interval (see Fig. S11C and Methods for details). Grey horizontal line indicates “no intervention”. (**D**) Counterfactual differential expression upon *KLF1* overexpression. Horizontal dashed line indicates –logFDR = 10. Vertical dashed lines indicate logFC = ±0.5. (**E**) Core network module explaining the counterfactual effect of *KLF1* overexpression. (**F**) GSEA cell type signature enrichments for the counterfactual expression change following CASCADE designed interventions for targeted differentiation towards 9 selected fetal cell types. Rows are fetal cell types used as design target state, followed by representative intervention designs in parentheses. Columns are MSigDB cell type signature gene sets. Dot sizes represent normalized enrichment score (NES), and colors represent −log10 false discovery rate (FDR). See Fig. S13 for the complete matrix of 62 cell types. (**G**) Amacrine cell marker expression observed in different treatment groups. Box plots indicate medians (centerlines), first and third quartiles (bounds of boxes) and 1.5× interquartile range (whiskers). (**H**) GSEA cell type signature enrichments for the observed expression change following CASCADE-designed *PAX6*+*TFAP2A* overexpression.

As a first application, CASCADE was trained on the K562 CRISPRa Perturb-seq dataset^17^ to design interventions for erythroid differentiation. Five erythroid single-cell datasets were used as the target state^32–35^ (Methods, Table S1). Among 32 experimentally perturbed candidate genes expressed at higher levels in erythroid than K562 cells, CASCADE unambiguously identified *KLF1* as the highest scoring gene (Fig. 4C) and the only gene above the 95% confidence interval-based cutoff (Fig. S11C, Methods), consistent with its established role as a key erythroid driver^36,37^. Counterfactual prediction of *KLF1* overexpression suggests a cell state shift in the erythroid direction (Fig. S11D), with up-regulation of various erythroid markers (Fig. 4D) attributable to a core *KLF1* network module (Fig. 4E).

Notably, the data-driven identification of *KLF1* was non-trivial, as it neither exhibits the highest differential expression in erythroid cells nor produced the most erythroid-like outcome as measured in the original Perturb-seq data (Fig. S11E). Other candidates such as *PRDM1* and *ZBTB1* produced more erythroid-like effects in the training data but were attributed to latent impact via *HBA2* or *KLF1* (Fig. S12), possibly due to the intrinsic differentiation tendency of K562 cells under stress^38,39^. Additionally, the CASCADE design suggests a higher *KLF1* overexpression rate (TPM ≈ 3,000) than in the original Perturb-seq experiment (TPM ≈ 300), which explains why simply matching measured perturbation effects would overlook important known biology, highlighting the need for a causality-oriented rational design approach.

To demonstrate the capability of CASCADE in a broader scope, we applied it to the human embryonic stem cell (hESC) TF overexpression atlas^40^, and used the inferred regulatory network to design combinatorial TF interventions that may drive differentiation towards all 62 well-defined fetal cell types in the human fetal gene expression atlas^41^ (Methods, Table S1). Counterfactual prediction of gene expression changes following the designed interventions demonstrated clear enrichment of cell type signatures for most targets, with relevant cell types activating shared functional programs (Fig. S13, also see Fig. 4F for cell types discussed below).

While for some cell types, the CASCADE designs overlap with factors identified in the original TF atlas study (Fig. 4F, *HNF4A* for intestinal epithelial cells and hepatocytes, *ERG* for vascular endothelial cells), the designed interventions diverge for more cell types. E.g., CASCADE identified *GATA1, KLF1* for erythroblast (Fig. S14A–D)^36,42^, the two most important erythroid drivers, compared to *KLF2* in the original study. Another example is metanephric cells, where CASCADE identified established nephric differentiation factors *PAX8, WT1* (Fig. S14E–H)^43,44^, compared to *CDX1* and *CDX2* in the original study^40^ which are better known as intestinal epithelial drivers instead^45^. Finally, CASCADE managed to identify known driver genes for many cell types that failed cell matching-based strategies used in the original study, such as *SPI1* for myeloid cells (Fig. S15A–D)^46^, *NR5A1* for adrenocortical cells (Fig. S15E–H)^47^, *KLF5, TP63* for squamous epithelial cells (Fig. S16A–D)^48,49^, *ISL1, PHOX2A* (Fig. S16E– H) for visceral neurons^50,51^, etc., highlighting CASCADE’s unique out-of-distribution generalization power.

As an effort to probe the limits of CASCADE’s rational intervention design, we sought to evaluate over highly specialized cell types that lacks well-established *in vitro* differentiation protocols. We focused on amacrine cells—retinal interneurons that refine and integrate visual signals within the inner plexiform layer. Upon jointly overexpressing the CASCADE-designed intervention factors *PAX6* and *TFAP2A* in hESC-derived induced neural progenitor cells (iNPCs), we successfully validated the strategy for inducing amacrine cells-like expression network (Fig. S17A–C) and observed robust up-regulation of multiple amacrine cell markers (Fig. 4G). Transcriptome-wide expression change caused by *PAX6*+*TFAP2A* overexpression also most prominently enriched for amacrine cell signature among closely related retinal neuronal subtypes, while depleting both NPC and ESC signatures (Fig. 4H, Table S12). Importantly, neither *PAX6* nor *TFAP2A* alone elicited a comparable induction of the amacrine program (Fig. 4G, Fig. S17D–F), highlighting the effectiveness of systematic multi-gene intervention designs.

## Discussion

Advances in genomic technologies provide increasingly rich and diverse experimental evidence for regulatory inference including both mechanistic regulatory footprints^20^ and perturbation outcomes^5^, collectively bringing us closer to the promise of creating comprehensive “virtual cells” capable of accurately simulating cellular behaviors *in silico* based on robust regulatory understandings^3^. Nonetheless, inconsistencies have been reported among different data types^52^, complicating the integration and translation of these heterogeneous data into coherent causal regulatory knowledge.

CASCADE addresses this challenge by harmonizing mechanistic regulatory knowledge with perturbation outcomes. Extending recent developments in DCD with a prior knowledge scaffold allows the model to focus its causal search over the vast genomic landscape and achieve efficient inference of causal GRNs encompassing thousands of genes, far exceeding the scope of canonical methods^7^, and enables interpretable counterfactual reasoning within a unified, causality-aware generative framework. Extensive benchmarking demonstrates that CASCADE consistently outperforms existing causal discovery and perturbation prediction methods across diverse synthetic and real-world datasets. Consequently, it is uniquely positioned to enable rational and quantitative intervention designs, e.g., to facilitate targeted cell programming with promising predictions that align well with established biological knowledge.

Apart from our causal GRN-based approach, models employing various neural network architectures including graph neural networks and transformer-based foundation models have also been proposed to predict the effects of unseen perturbations^24–26,29,53^, and conversely, to identify perturbations that would yield desired responses^28^. However, these methods rely on statistical associations encoded within latent cell and gene embeddings that do not necessarily imply causal understanding, which could be a major reason behind their inferior performance over linear baselines^31^. Consistently, our empirical assessments well demonstrated that explicitly modeling the causal GRN in the original gene space not only enhances interpretability in terms of mechanistic understanding, but also improves the generalization power of perturbation predictions, which is crucial for practically useful intervention design.

CASCADE’s Bayesian framework provides uncertainty estimates for individual regulatory relations, which can potentially be exploited to highlight ambiguous network segments and guide active learning strategies to iteratively refine network inference^54^. Of note, in efforts to model regulatory effects not captured by current transcriptomic measurements like chromatin remodeling, post-translational modifications, and subcellular localizations^55^, an external-encoded latent variable is explicitly introduced into current CASCADE. While we recognize that such “hybrid design” does not necessarily hold theoretical guarantee on causal identifiability, empirical evaluation well confirmation its effectiveness and generalizability (Fig. S5), and may enable a progressive path where further information could be gradually articulated as part of the causal graph as our understanding of regulatory biology and measurement technologies continue to evolve.

We recognize that our assumption of an acyclic GRN structure under a causal framework contrasts with the prevalence of feedback loops in biological systems. Typically, cyclic networks are interpreted as dynamic systems over the temporal dimension, a scenario challenging to resolve from non-temporal data alone^56^. Our Bayesian formulation partially alleviates this issue by permitting segments of feedback loops to reside in distinct SVGD particles (Fig. S6C, D, Fig. S7C). Nevertheless, more rigorous causal modeling of feedback loops demands additional theoretical considerations, such as the steady state assumption^57^, to ensure accurate causal inference.

Currently, CASCADE employs MLPs as structural equations within the causal network. While effective, recent advances in neural network architectures, including transformer models, have demonstrated exceptional performance across various domains^58^. Unlike natural language processing, which benefits from the inherently causal sequential structure of texts^59^, tabular gene expression data lack such causal ordering. Integrating adaptive causal learning strategies in CASCADE with powerful transformer architectures may enhance the capability of current single-cell foundation models, particularly for out-of-distribution tasks such as perturbation predictions.

Beyond genetic perturbations, large-scale chemical perturbation screens are also gaining prominence^60^. Although CASCADE currently lacks direct chemical modeling capabilities, it holds the potential to elucidate immediate gene targets responsive to these perturbations, either by incorporating chemical features into latent representations during training, or by solving intervention design problems to identify genetic perturbations that recapitulate chemical-induced effects.

In conclusion, CASCADE presents a significant step towards integrating diverse regulatory evidence within a unified causality-aware framework. We envision CASCADE playing an enduring role in the ongoing endeavor to decipher and leverage the complete causal regulatory network to engineer cellular behaviors. The whole package of CASCADE is available online at https://github.com/gao-lab/CASCADE.

## Acknowledgments

The authors would like to thank Drs. Fuchou Tang, Xiaoliang Sunney Xie, Yiqin Gao, Zemin Zhang, Letian Tao, Cheng Li, Jian Lu, Hsiang-Ying Lee, Nan Liang, Shunkang Wu (at Peking University), Fan Zhou (at Tsinghua University) and Yang Ding (at Changping Laboratory) for their helpful discussions and comments during the study, as well as authors of the datasets used in this work for their kindly help.

This work was supported by funds from the National Science and Technology Major Project (2022ZD0115004), China Postdoctoral Science Foundation (grant no. 2023T160009), the State Key Laboratory of Protein and Plant Gene Research and the Beijing Advanced Innovation Center for Genomics (ICG) at Peking University, the CAS Project for Young Scientists in Basic Research (grant no. YSBR-012), as well as the Changping Laboratory.

Part of the analysis was performed on the Computing Platform of the Center for Life Sciences of Peking University and supported by the High-performance Computing Platform of Peking University.

## Author contributions

G.G. conceived the study and supervised the research; Z.J.C. designed and implemented the computational framework; Z.J.C. and C.G. compiled the scaffold graphs; Z.J.C. and C.L. conducted benchmarks and case studies, with guidance from G.G.; F.Z and T.G. conducted hESC differentiation validation experiments, with guidance from F.G.; Z.J.C. and P.L. analyzed the experiment data; Z.J.C. and G.G. wrote the manuscript, with inputs from all authors.

## Supplementary figures and tables

**Fig. S1.**
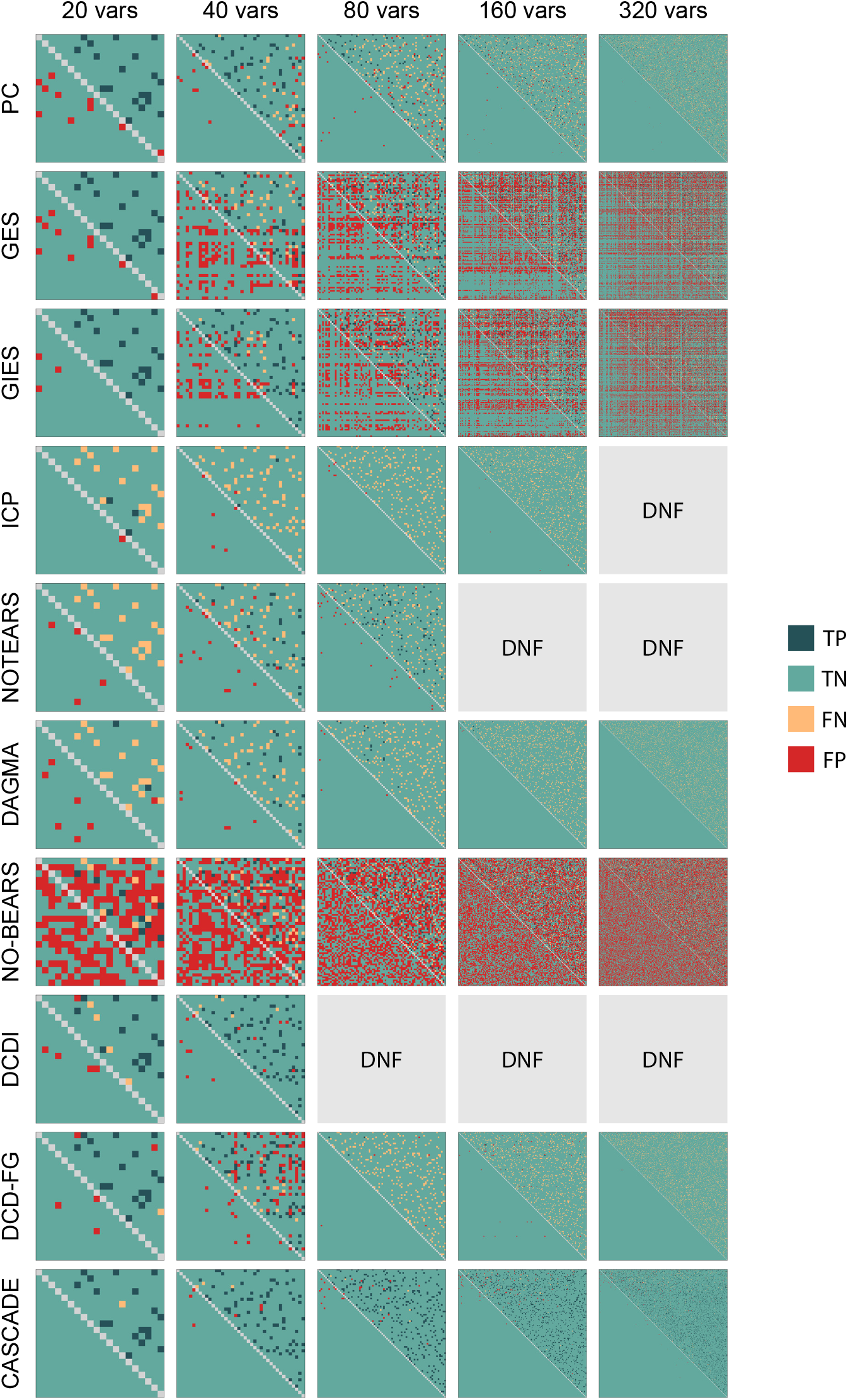
Example adjacency graphs inferred by different causal discovery methods across different numbers of variables. TP, true positive; TN, true negative; FN, false negative; FP, false positive; DNF, did not finish.

**Fig. S2.**
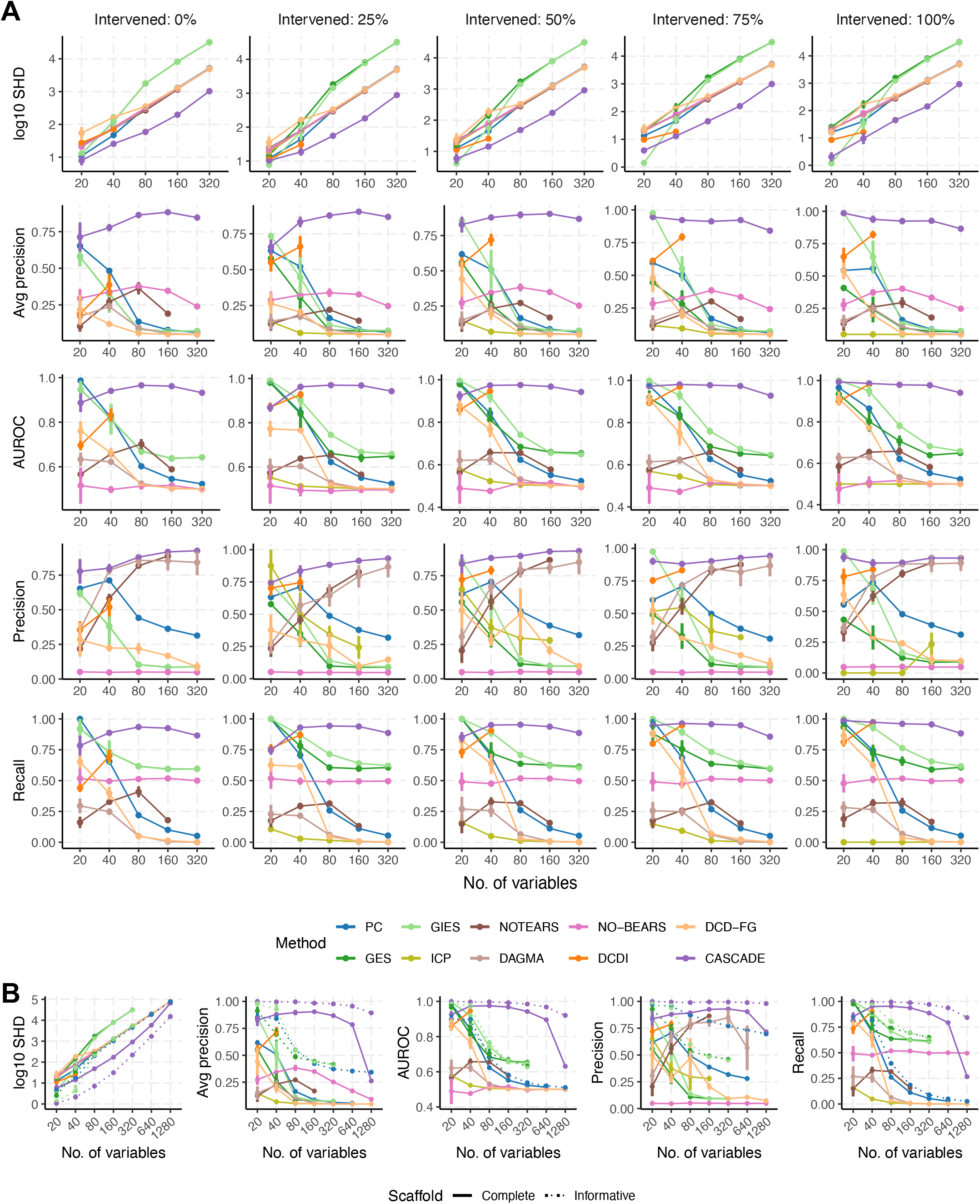
Extended comparison of causal discovery accuracies with simulated data. (**A**) Accuracy of different causal discovery methods across different numbers of variables and intervention fractions, as quantified by log_10_ structural Hamming distance (SHD), average precision, area under receiver operating characteristic (AUROC), precision and recall. N = 4 with different ground truth DAGs. Error bars indicate mean ± s.d. (**B**) Accuracy of different causal discovery methods with up to 1,280 variables and 50% intervened fraction. The informative scaffold was obtained by corrupting the true DAG with 10% false positive edges. N = 4 with different true DAGs. N = 4 with different ground truth DAGs.

**Fig. S3.**
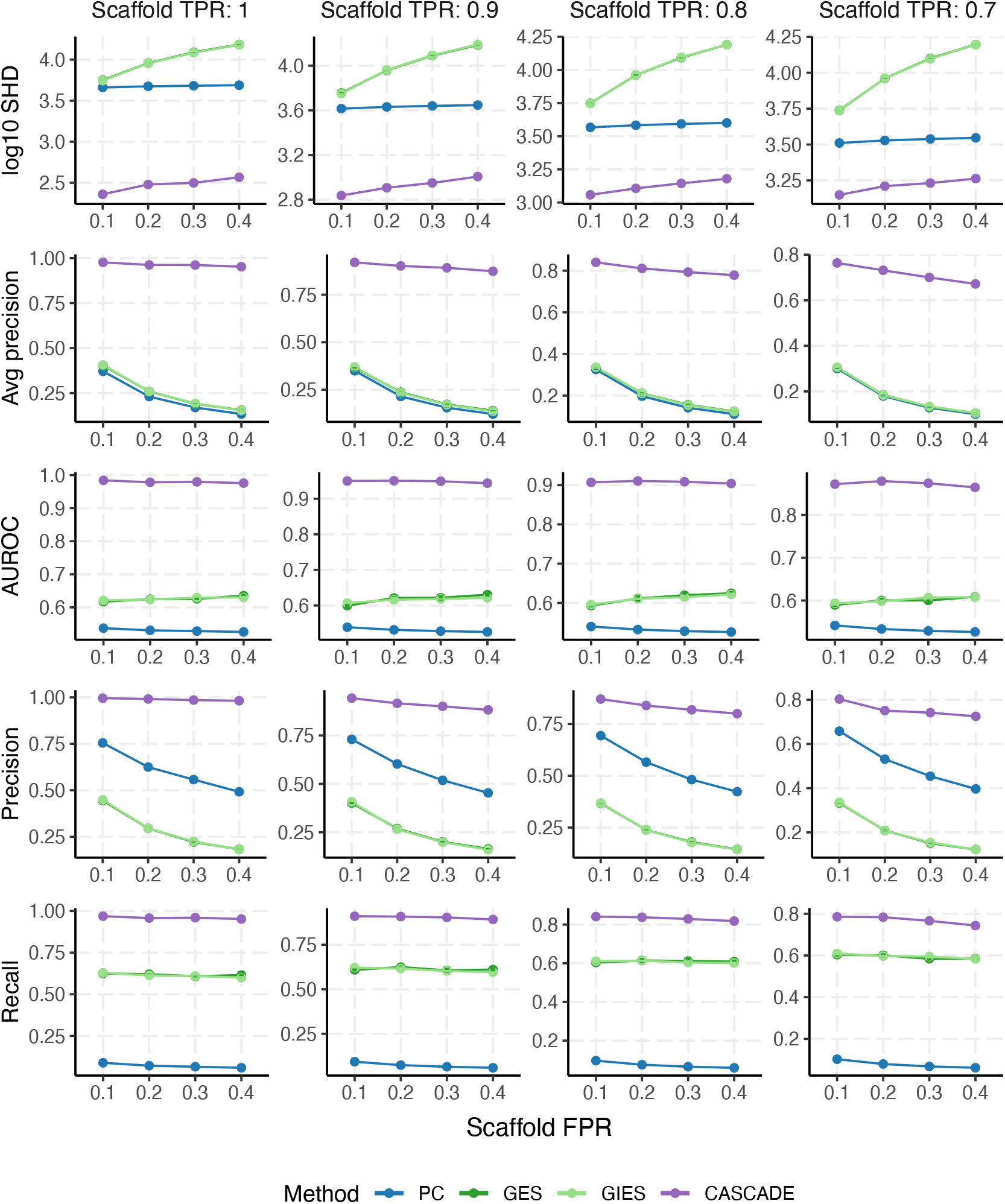
Causal discovery accuracy across different TPRs and FPRs in the scaffold graph. N = 4 with different corrupted scaffold graphs. Error bars indicate mean ± s.d.

**Fig. S4.**
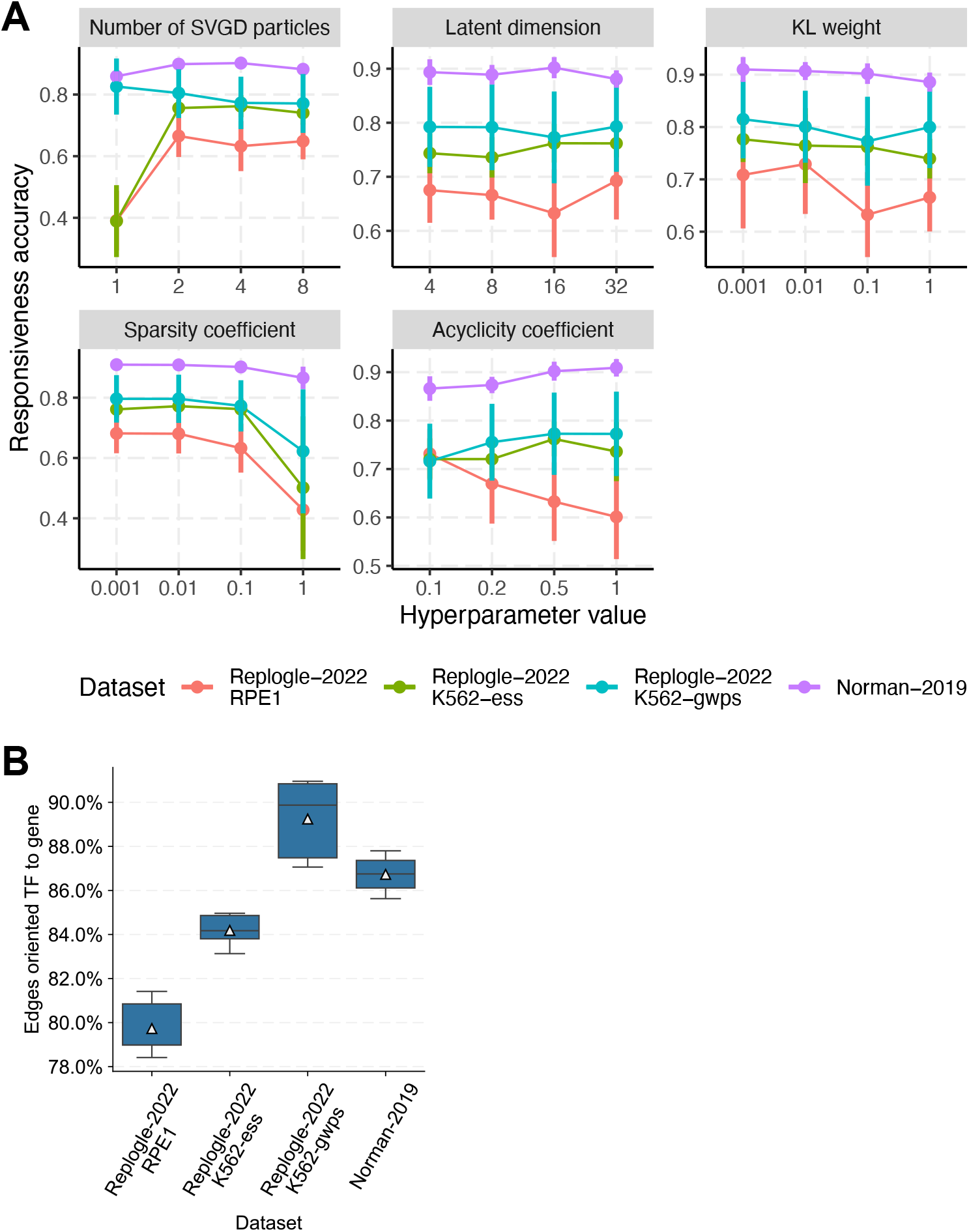
Hyperparameter robustness and edge orientation accuracy of CASCADE in Perturb-seq causal discovery. (**A**) Hyperparameter robustness. N = 5 with different train/test splits. The hyperparameters are denoted in the Methods section as *n* for number of SVGD particles, *Z* for latent dimension, *β* for latent KL weight, *η*_*λ*_ for sparsity coefficient, and *η*_*α*_ for acyclicity coefficient. Error bars indicate mean ± s.d. (**B**) Edge orientation accuracy among undirected scaffold edges involving one TF, assuming the correct orientation points outward from TFs. N = 5 with different train/test splits. Box plots indicate medians (centerlines), mean (triangles), first and third quartiles (bounds of boxes) and 1.5× interquartile range (whiskers).

**Fig. S5.**
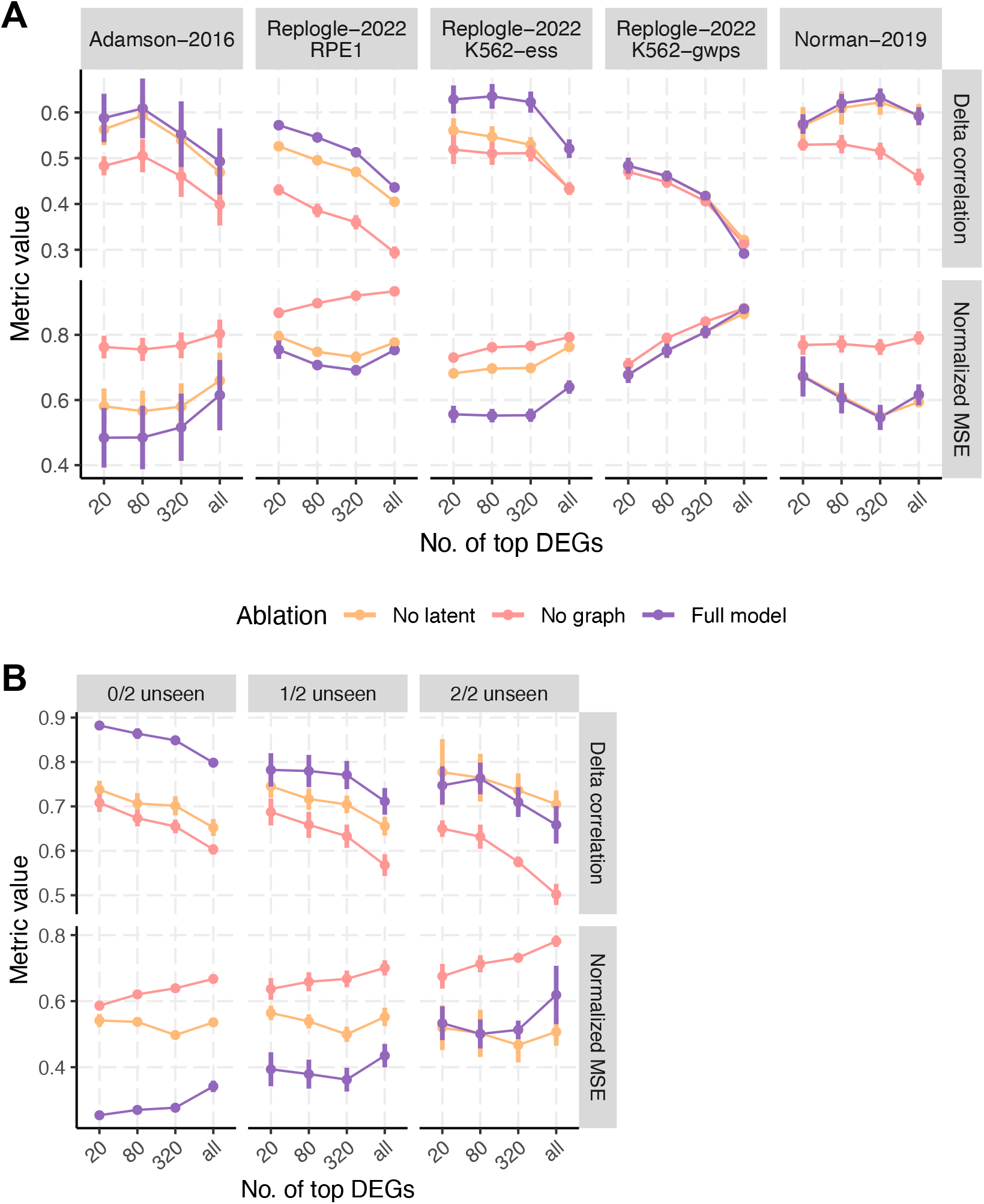
Ablation experiment for CASCADE perturbation prediction. Prediction accuracy with either the full model, an ablated model without latent factors, or an ablated model without causal graph propagation in (**A**) single gene perturbation prediction, and (**B**) double gene perturbation prediction.

**Fig. S6.**
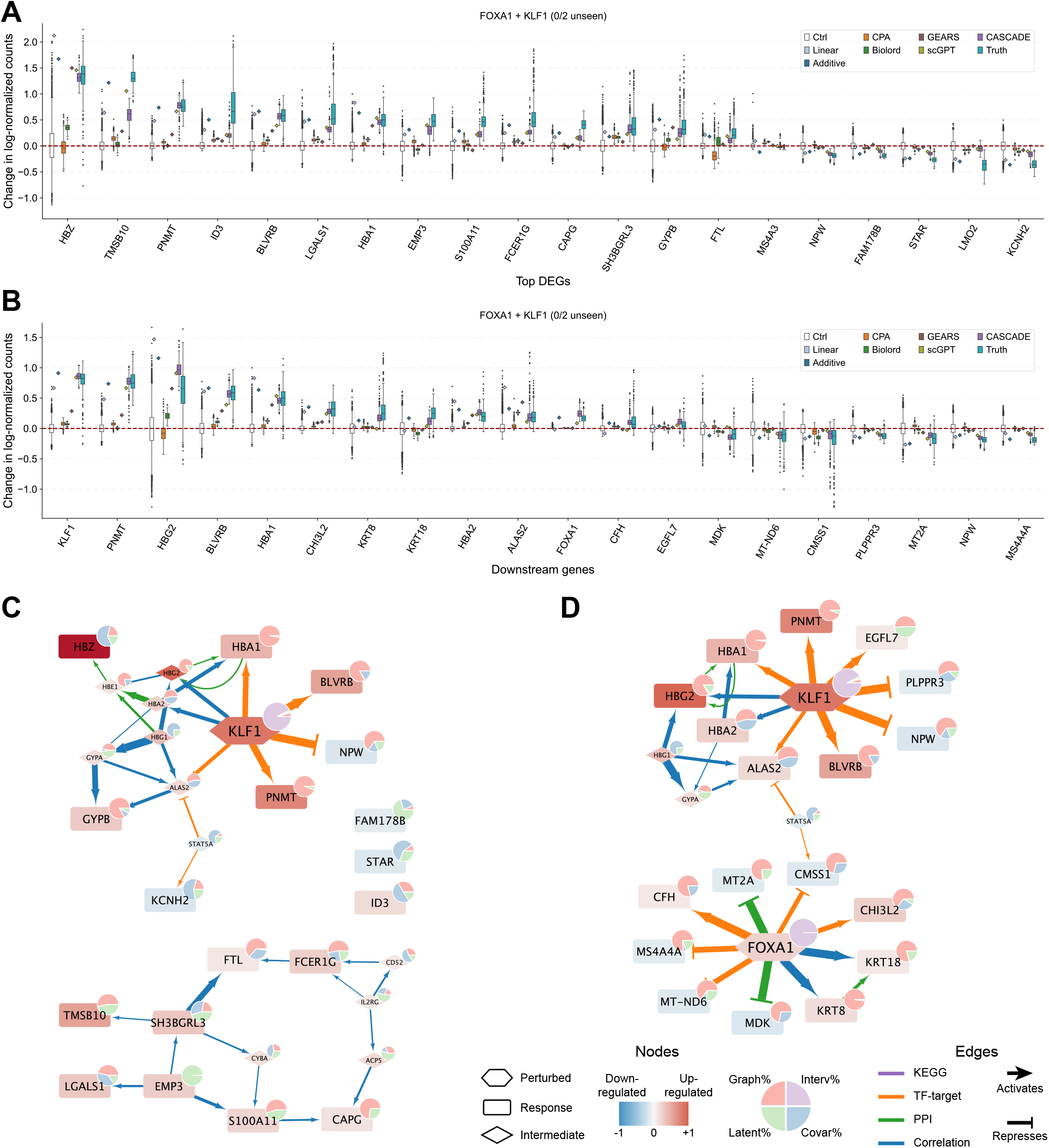
Perturbation prediction and interpretation for FOXA1+KLF1 activation. (**A**) Prediction comparison for the top 20 DEGs upon FOXA1 and KLF1 activation. (**B**) Prediction comparison for the model inferred downstream genes of FOXA1 and KLF1. Box plots indicate medians (centerlines), first and third quartiles (bounds of boxes) and 1.5× interquartile range (whiskers). (**C**) Prediction interpretation for the top 20 DEGs. (**D**) Prediction interpretation for downstream genes. Node colors indicate predicted expression changes. Pie charts indicate contribution proportions. Edge colors indicate scaffold evidence types. Edge widths indicate contribution amount.

**Fig. S7.**
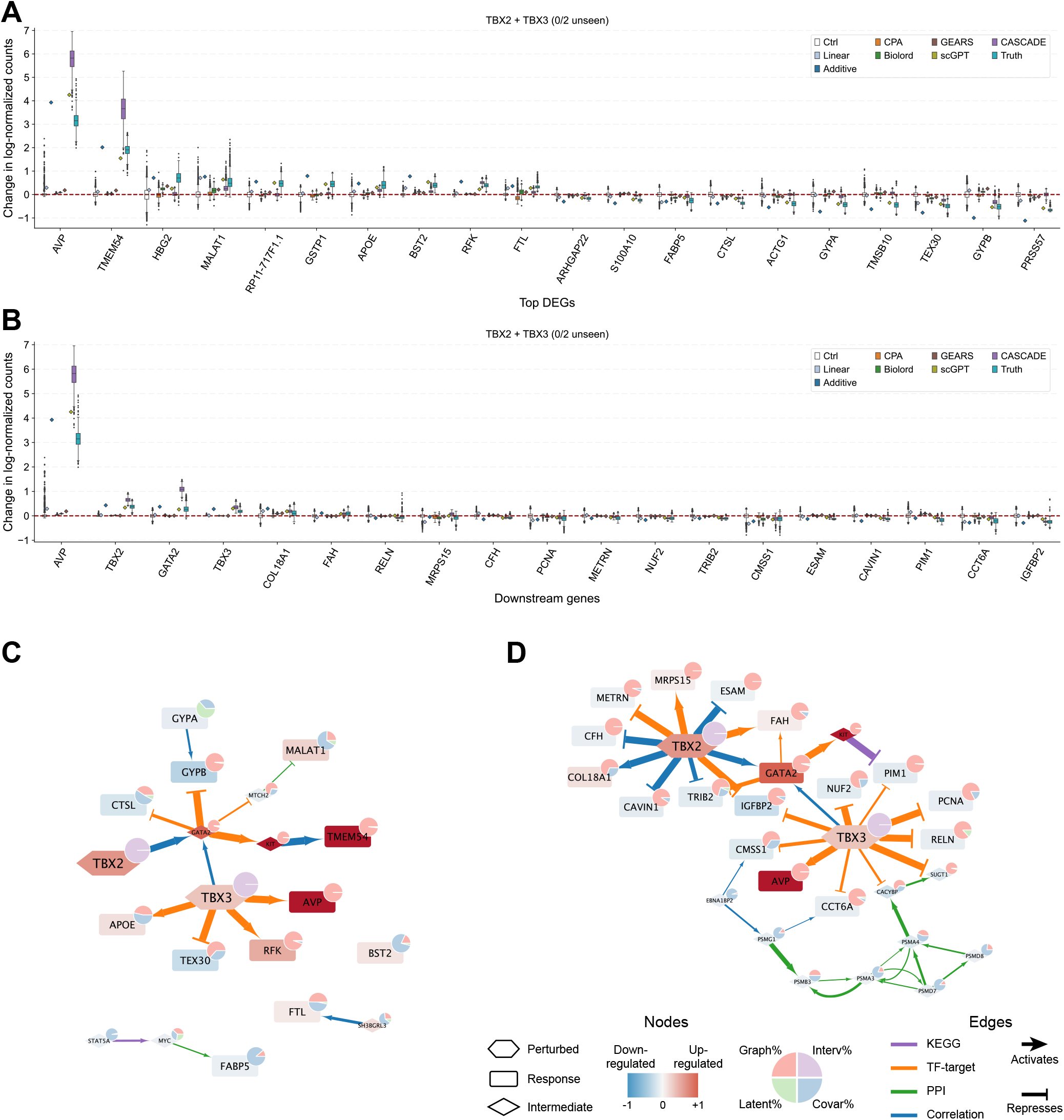
Perturbation prediction and interpretation for TBX2+TBX3 activation. (**A**) Prediction comparison for the top 20 DEGs upon TBX2 and TBX3 activation. (**B**) Prediction comparison for the model inferred downstream genes of TBX2 and TBX3. Box plots indicate medians (centerlines), first and third quartiles (bounds of boxes) and 1.5× interquartile range (whiskers). (**C**) Prediction interpretation for the top 20 DEGs. (**D**) Prediction interpretation for downstream genes. Node colors indicate predicted expression changes. Pie charts indicate contribution proportions. Edge colors indicate scaffold evidence types. Edge widths indicate contribution amount.

**Fig. S8.**
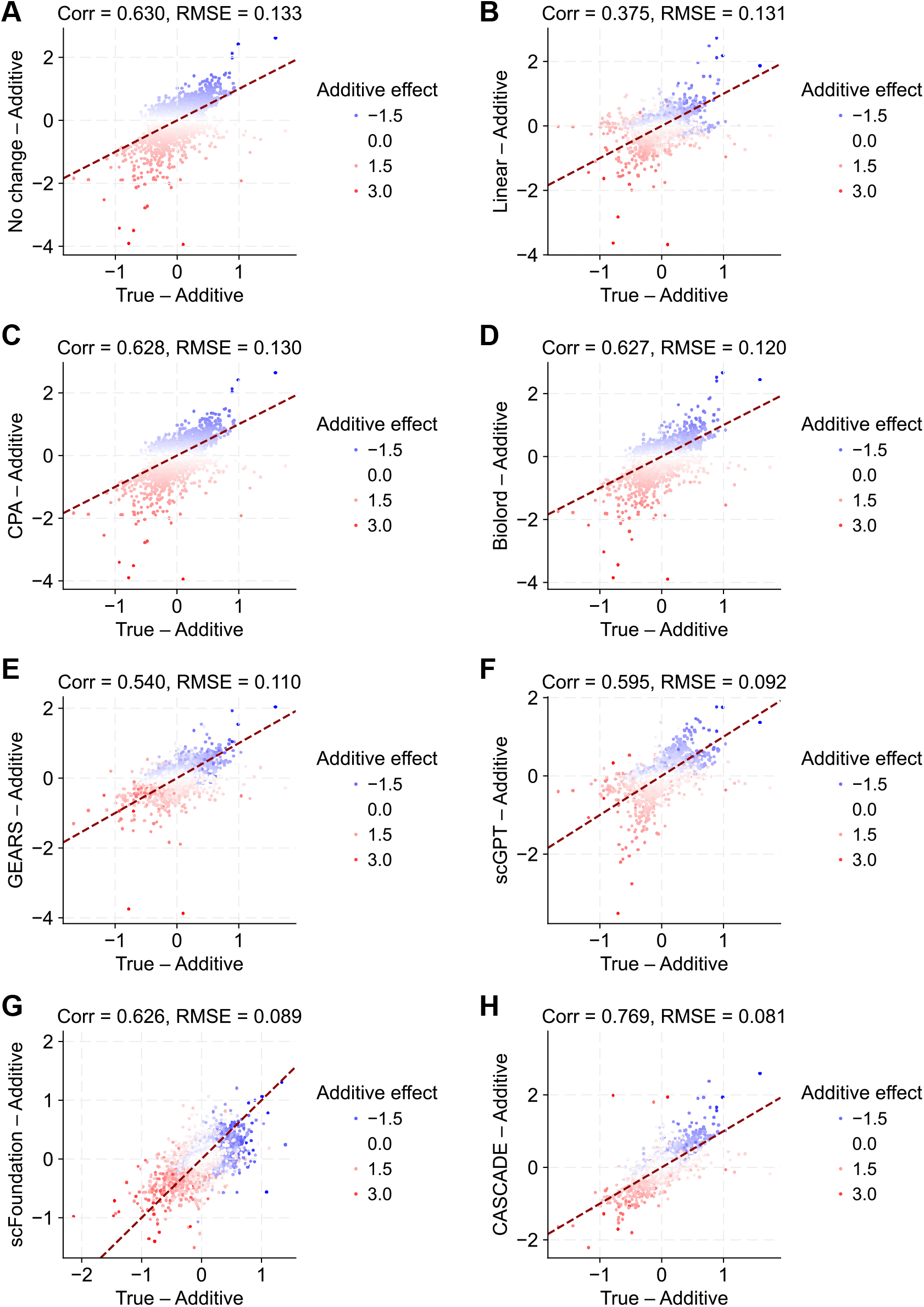
Prediction accuracy for non-additive residual effects in double-gene perturbations. Predicted non-additive effect vs true non-additive effect for (**A**) no change (predicting control), (**B**) linear baseline, (**C**) CPA, (**D**) Biolord, (**E**) GEARS, (**F**) scGPT, (**G**) scFoundation, (**H**) CASCADE. Genes are colored by additive effect vs control.

**Fig. S9.**
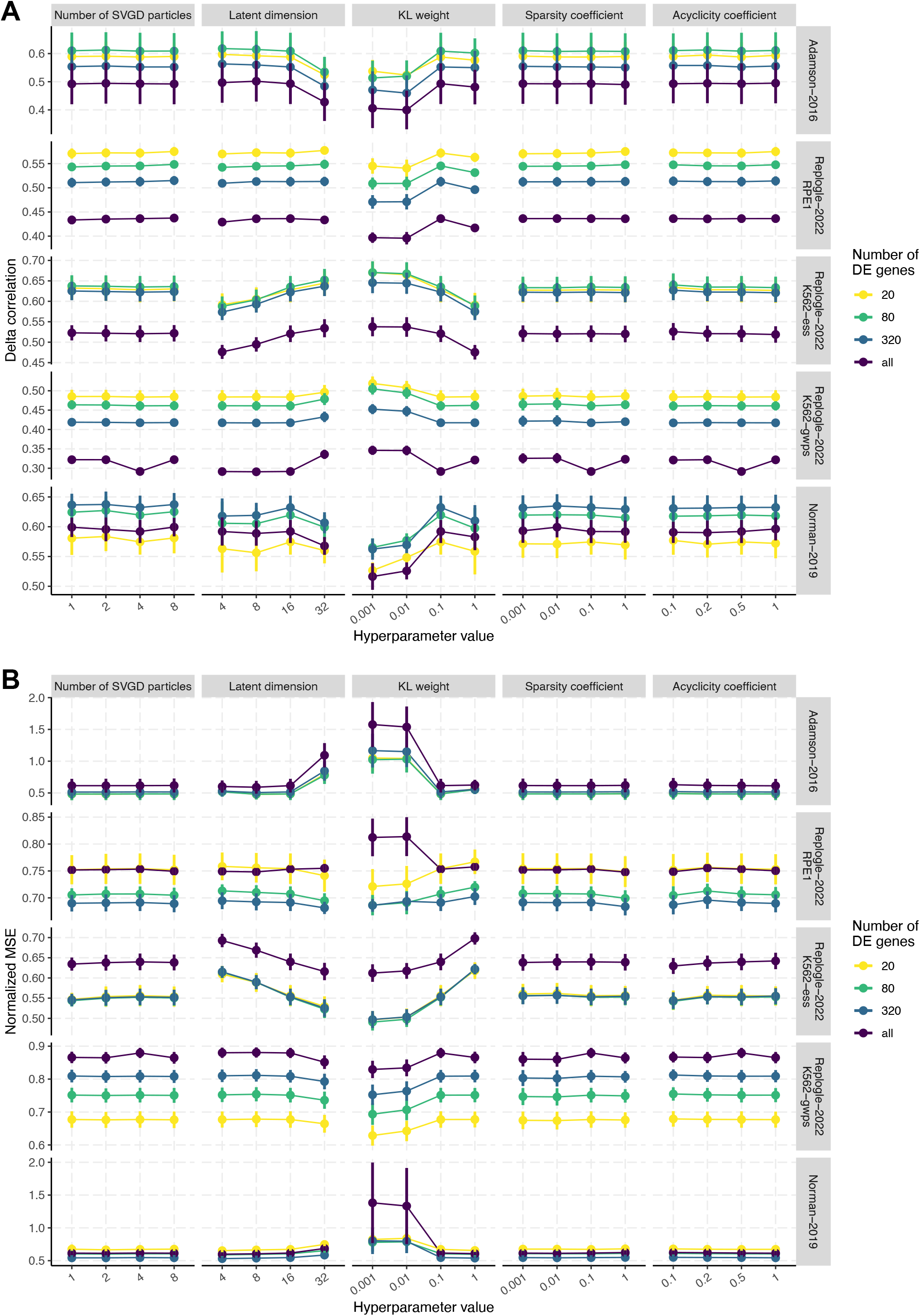
Hyperparameter robustness of CASCADE in single-gene perturbation prediction. Prediction accuracy was quantified by (**A**) delta correlation, and (**B**) normalized MSE. N = 5 with different train/test splits. The hyperparameters are denoted in the Methods section as *n* for number of SVGD particles, *Z* for latent dimension, *β* for latent KL weight, *η*_*λ*_ for sparsity coefficient, and *η*_*α*_ for acyclicity coefficient. Error bars indicate mean ± s.d.

**Fig. S10.**
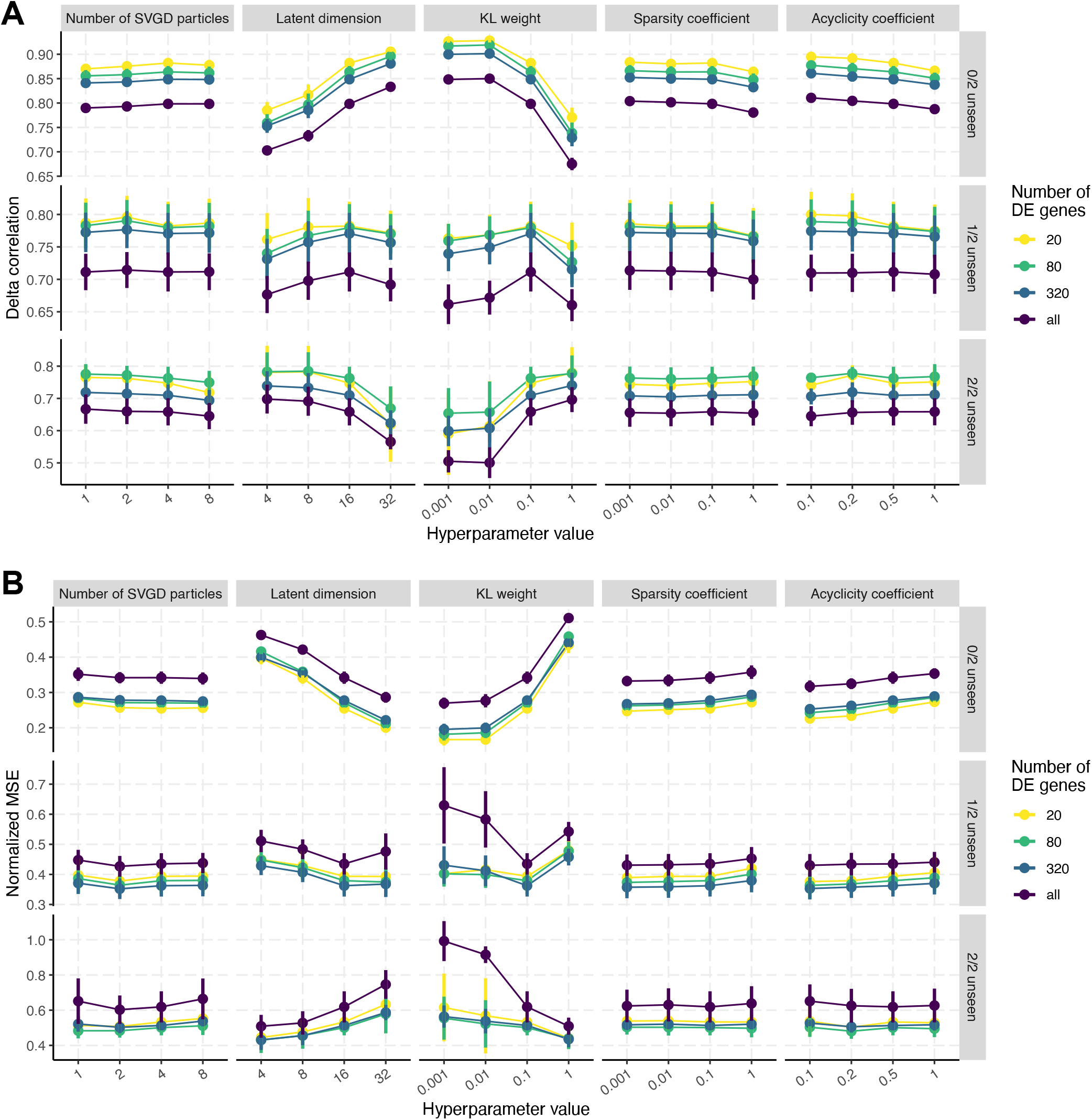
Hyperparameter robustness of CASCADE in double-gene perturbation prediction. Prediction accuracy was quantified by (**A**) delta correlation, and (**B**) normalized MSE. N = 5 with different train/test splits. The hyperparameters are denoted in the Methods section as *n* for number of SVGD particles, *Z* for latent dimension, *β* for latent KL weight, *η*_*λ*_ for sparsity coefficient, and *η*_*α*_ for acyclicity coefficient. Error bars indicate mean ± s.d.

**Fig. S11.**
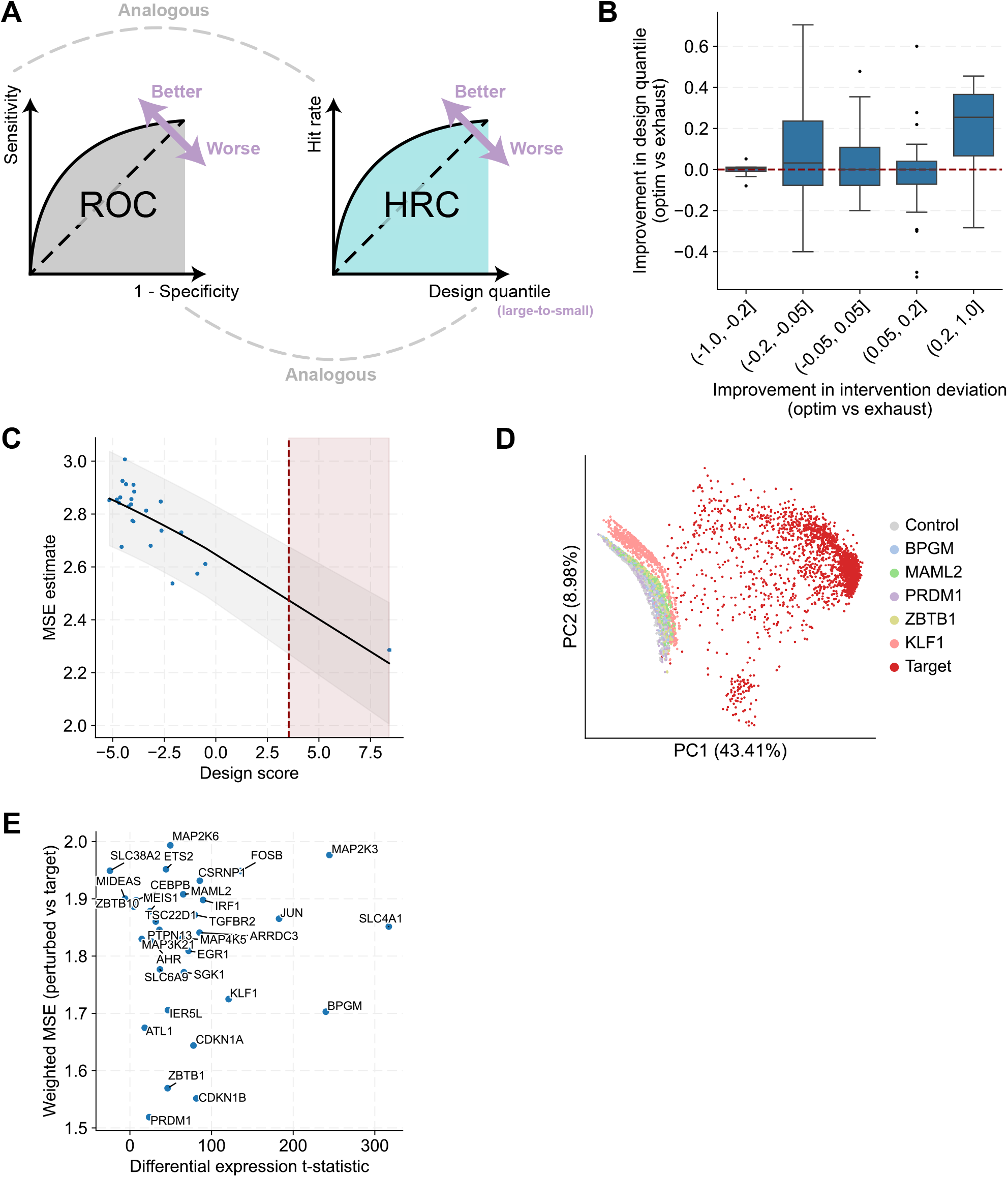
Advantage of optimization-based design and application in erythroid differentiation. (**A**) Illustration of the hit rate curve (HRC). (**B**) Comparison between optimization-based vs exhaustive search-based CASCADE design. The x-axis represents improvement in deviation from the true interventional magnitudes of the intervened genes themselves, achieved by the optimization-based approach (where interventional magnitudes are optimizable) vs exhaustive search (where interventional magnitudes are fixed as the training mean). The y-axis represents improvement in the quantile of true interventions among design rankings. Box plots indicate medians (centerlines), first and third quartiles (bounds of boxes) and 1.5× interquartile range (whiskers). (**C**) Design scores vs MSEs between counterfactual outcome and the target state (see Methods for details). Black solid line and grey strip indicate smoothed mean and 95% confidence interval by Gaussian process regression. Red vertical line indicates the empirical design score cutoff covering the smallest MSE within 95% confidence interval. (**D**) Joint PCA visualization of erythroid cells and counterfactual K562 cells under different interventions. (**E**) Landscape of all candidate genes, with the x-axis indicating differential expression of the candidate genes themselves between erythroid cells and K562 cells, and the y-axis indicating the transcriptome-wide weighted MSE between erythroid cells and measured perturbation outcomes.

**Fig. S12.**
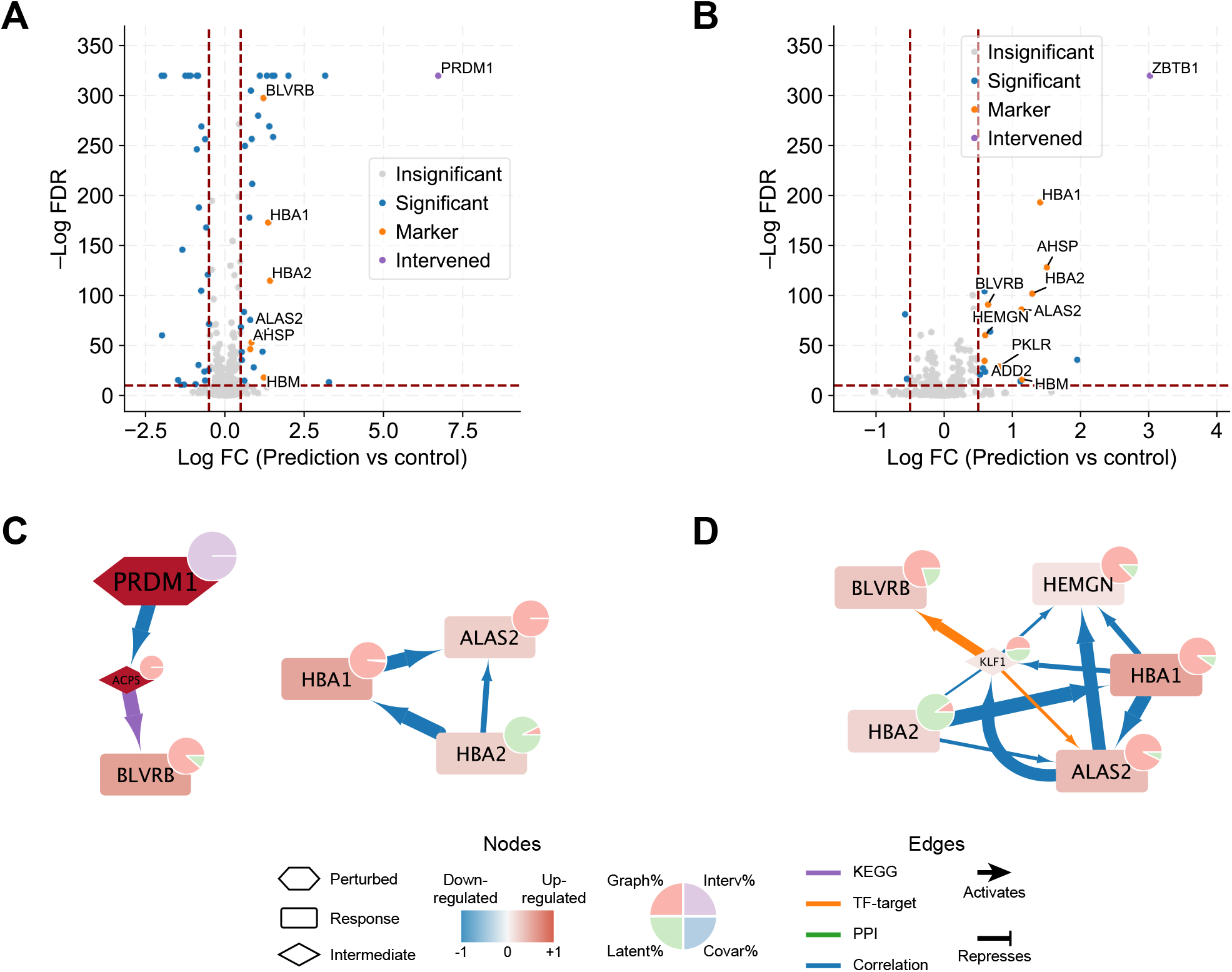
Intervention design for K562-to-erythroid differentiation. (**A**) Counterfactual differential expression upon PRDM1 overexpression. (**B**) Counterfactual differential expression upon ZBTB1 overexpression. Horizontal dashed line indicates –logFDR = 10. Vertical dashed lines indicate logFC = ±0.5. (**C**) Core network module explaining the counterfactual effect of PRDM1 overexpression. (**D**) Core network module explaining the counterfactual effect of ZBTB1 overexpression.

**Fig. S13.**
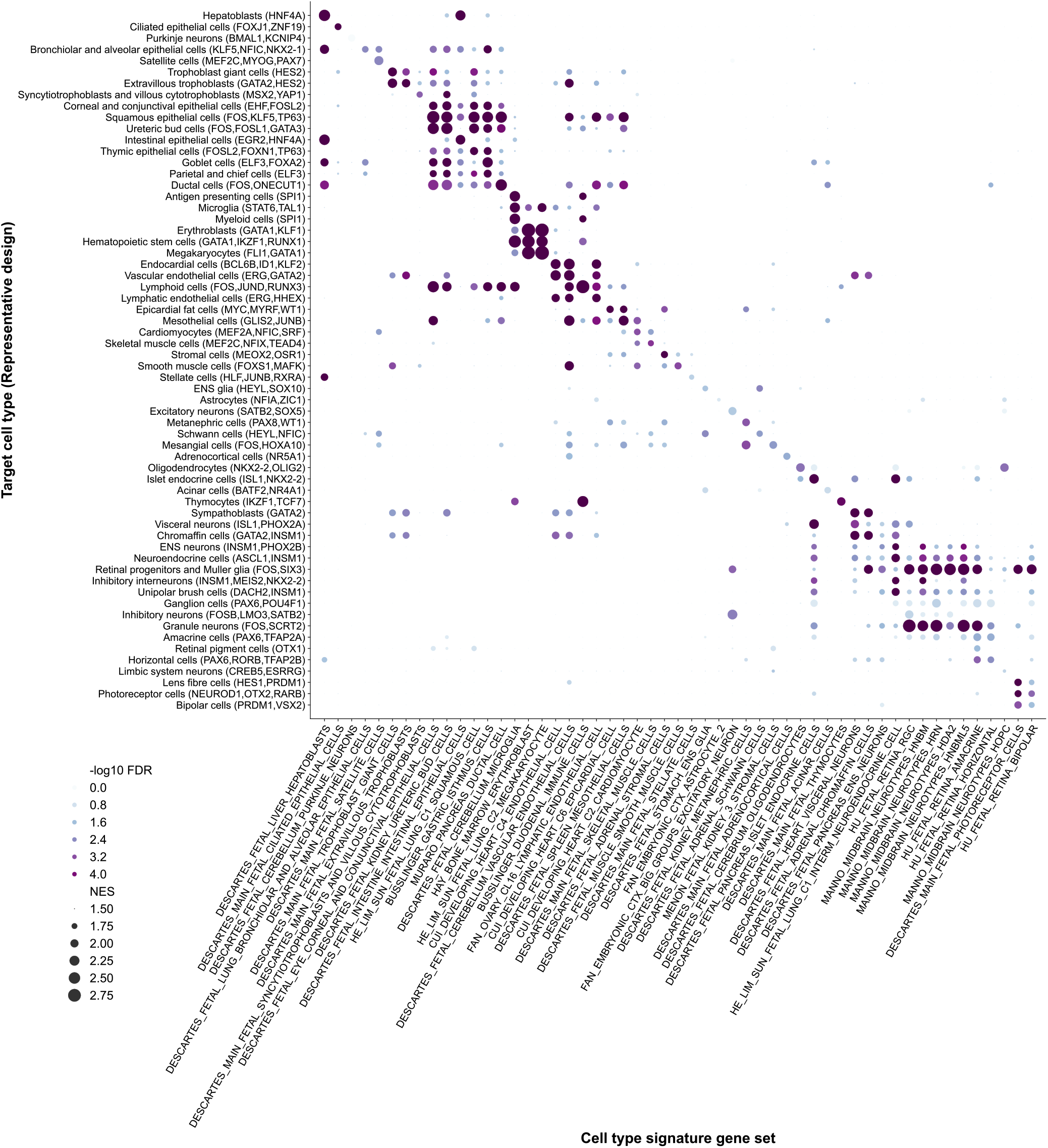
Rationally designed hESC differentiation towards 62 distinct fetal cell types. GSEA cell type signature enrichments for the counterfactual expression change following CASCADE designed interventions for targeted differentiation towards different fetal cell types. Rows are fetal cell types used as design target state, followed by representative intervention designs in parentheses. Columns are MSigDB cell type signature gene sets. Dot sizes represent normalized enrichment score (NES), and colors represent −log10 false discovery rate (FDR).

**Fig. S14.**
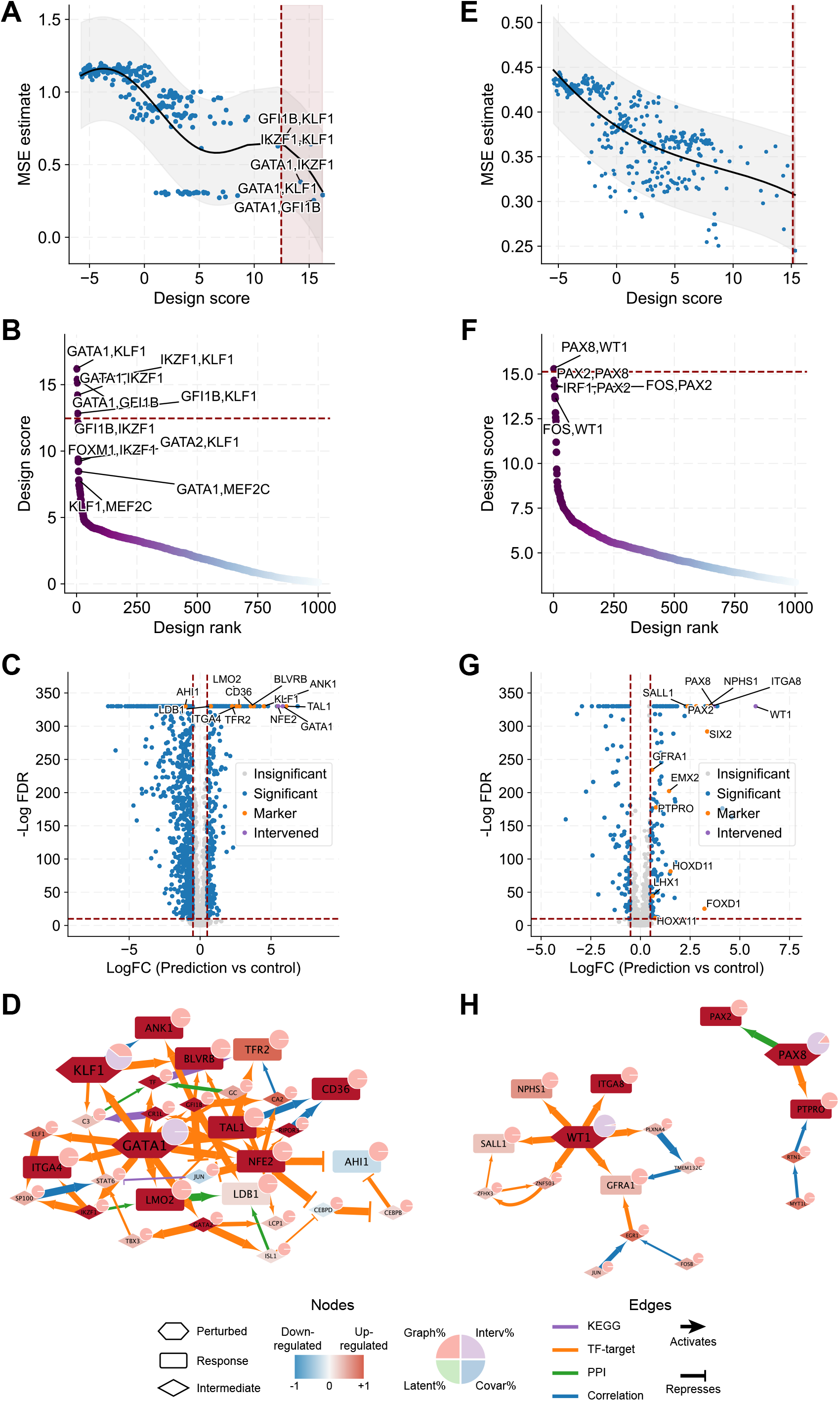
Intervention design for hESC-to-erythroblast and metanephric cell differentiation. (**A**–**D**) Intervention design for hESC-to-erythroblast differentiation, with (**A**) design scores vs MSEs between counterfactual outcomes and erythroblasts, (**B**) top ranked interventions, (**C**) counterfactual differential expression after *GATA1, KLF1* overexpression, (**D**) core network module explaining the counterfactual effect of *GATA1, KLF1* overexpression. (**E**–**H**) Intervention design for hESC-to-metanephric cell differentiation, with (**E**) design scores vs MSEs between counterfactual outcomes and metanephric cells, (**F**) top ranked interventions, (**G**) counterfactual differential expression after *PAX8, WT1* overexpression, (**H**) core network module explaining the counterfactual effect of *PAX8, WT1* overexpression.

**Fig. S15.**
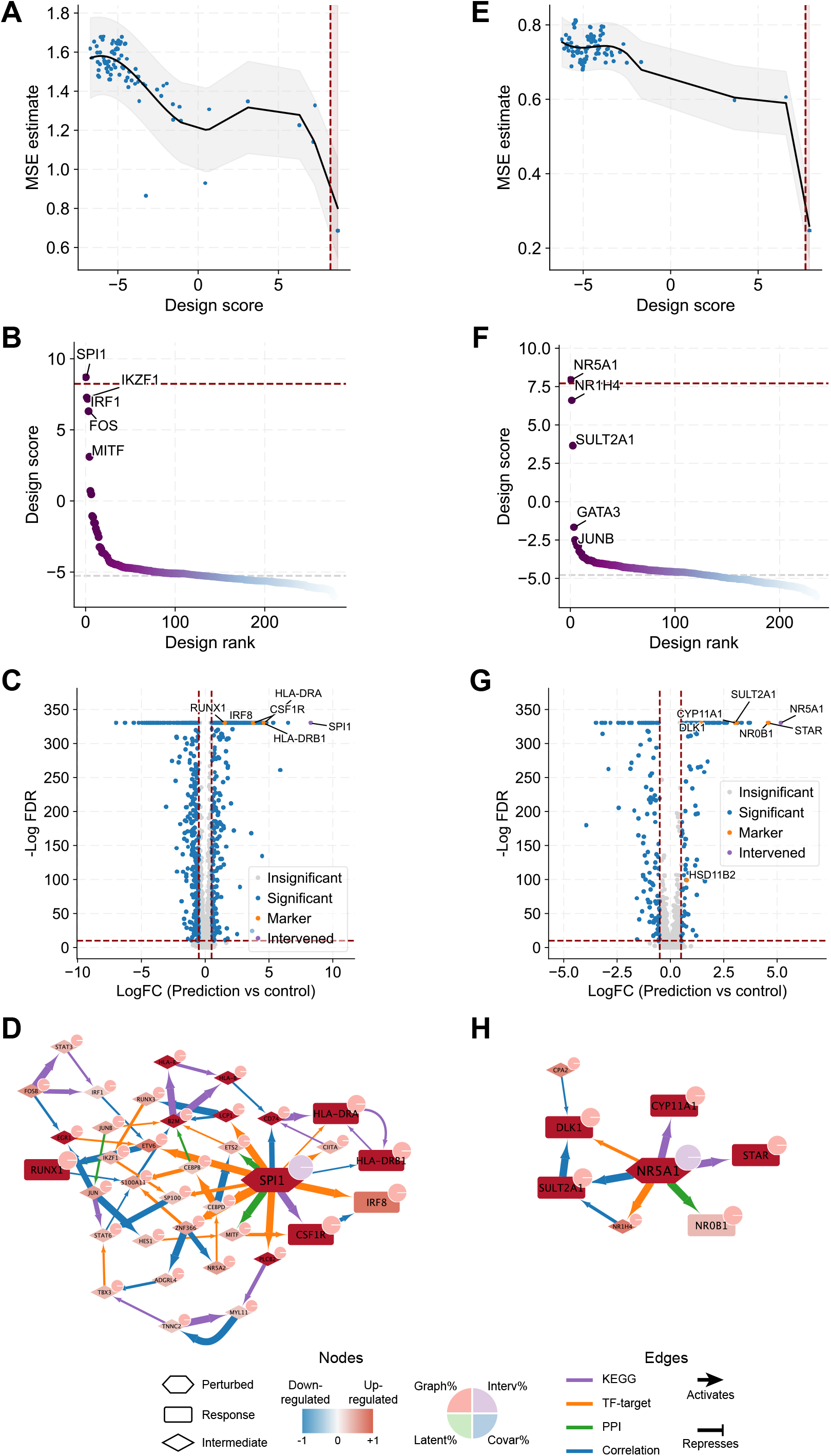
Intervention design for hESC-to-myeloid and adrenocortical cell differentiation. (**A**–**D**) Intervention design for hESC-to-myeloid cell differentiation, with (**A**) design scores vs MSEs between counterfactual outcomes and myeloid cells, (**B**) top ranked interventions, (**C**) counterfactual differential expression after *SPI1* overexpression, (**D**) core network module explaining the counterfactual effect of *SPI1* overexpression. (**E**–**H**) Intervention design for hESC-to-adrenocortical cell differentiation, with (**E**) design scores vs MSEs between counterfactual outcomes and adrenocortical cells, (**F**) top ranked interventions, (**G**) counterfactual differential expression after *NR5A1* overexpression, (**H**) core network module explaining the counterfactual effect of *NR5A1* overexpression.

**Fig. S16.**
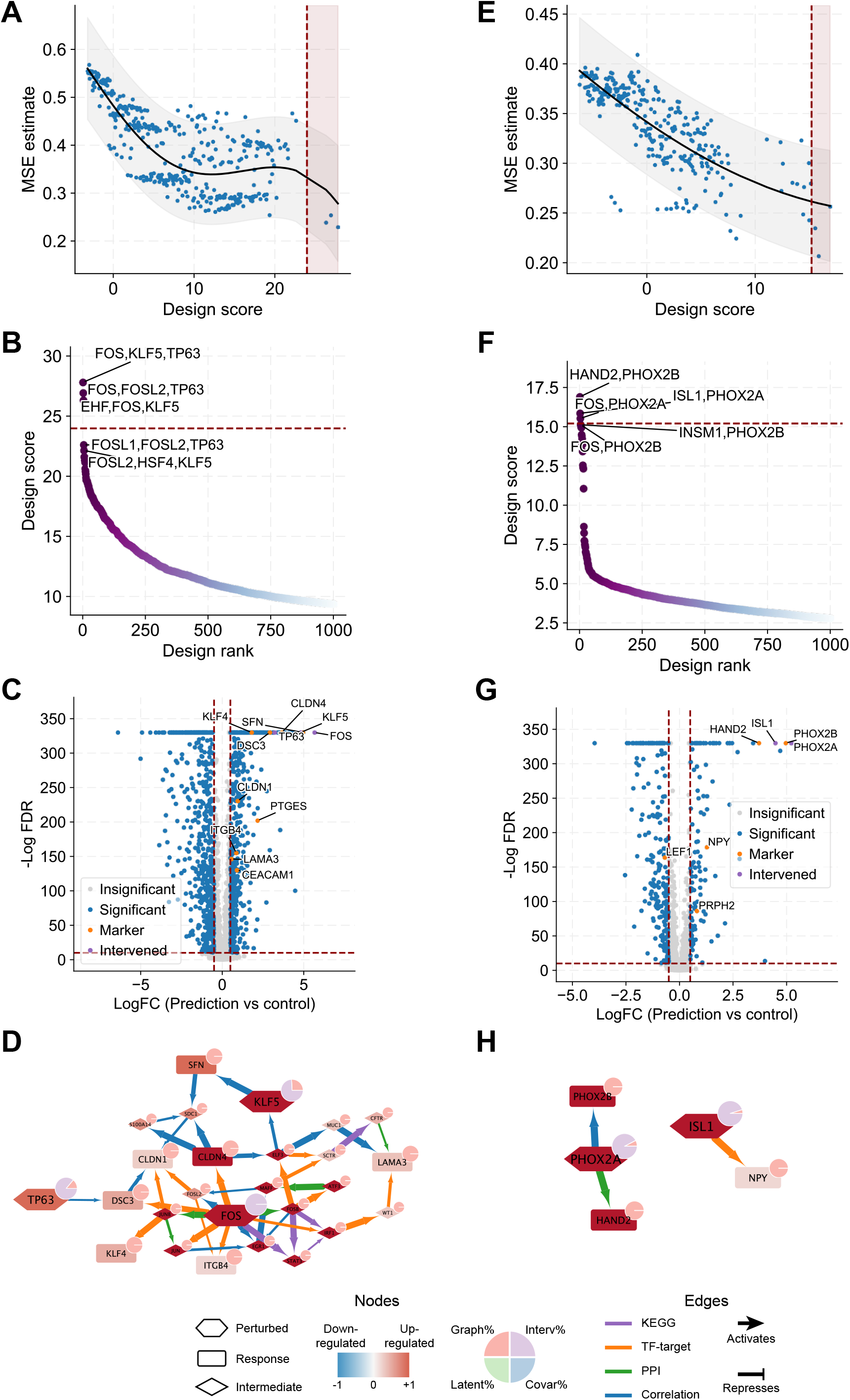
Intervention design for hESC-to-squamous epithelial cell and visceral neuron differentiation. (**A**–**D**) Intervention design for hESC-to-squamous epithelial cell differentiation, with (**A**) design scores vs MSEs between counterfactual outcomes and squamous epithelial cells, (**B**) top ranked interventions, (**C**) counterfactual differential expression after *FOS, KLF5, TP63* overexpression, (**D**) core network module explaining the counterfactual effect of *FOS, KLF5, TP63* overexpression. (**E**–**H**) Intervention design for hESC-to-visceral neuron differentiation, with (**E**) design scores vs MSEs between counterfactual outcomes and visceral neurons, (**F**) top ranked interventions, (**G**) counterfactual differential expression after *ISL1, PHOX2A* overexpression, (**H**) core network module explaining the counterfactual effect of *ISL1, PHOX2A* overexpression.

**Fig. S17.**
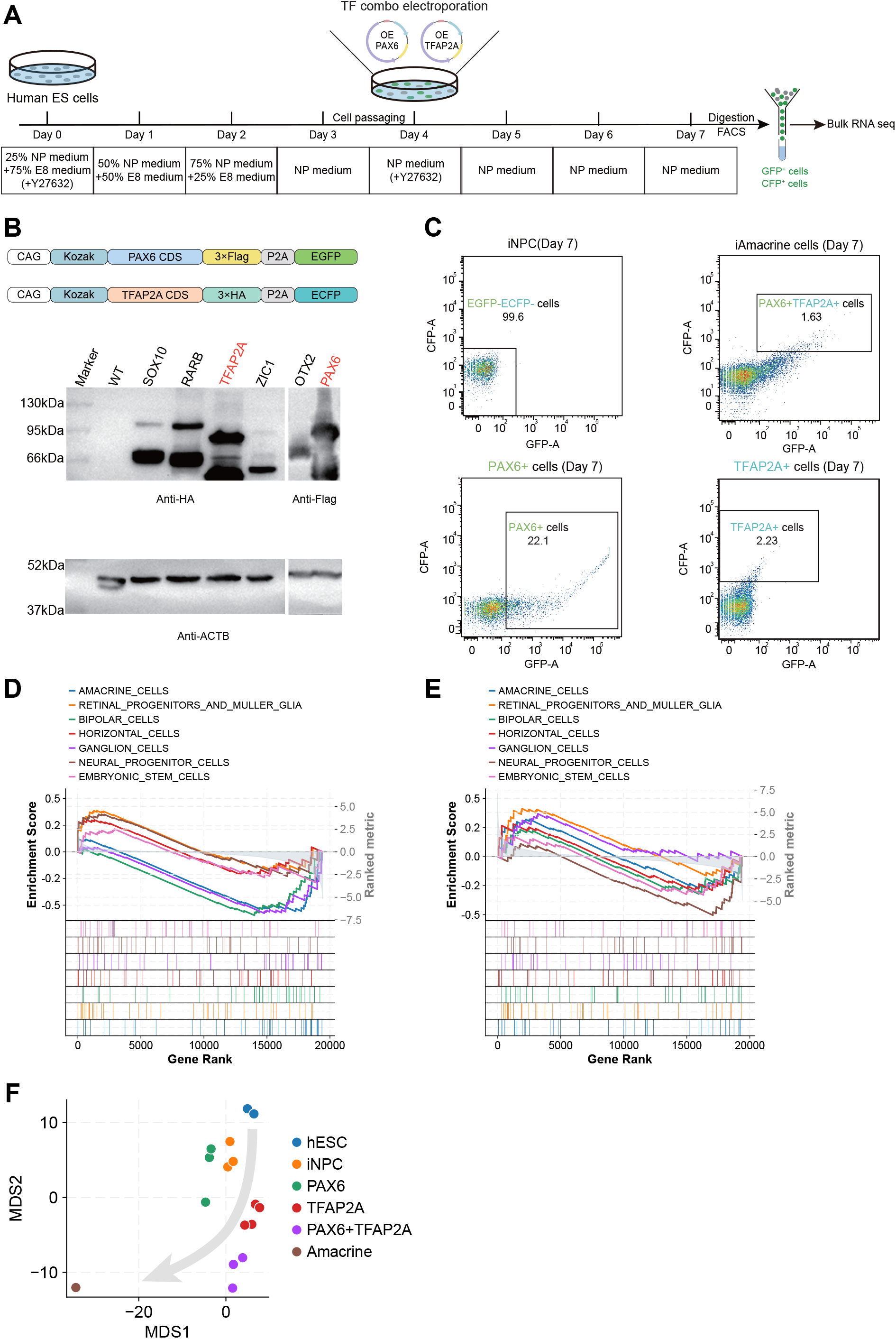
Stepwise in vitro transcription factor-guided differentiation of human embryonic stem cells into amacrine cell-like expression networks. (**A**) Experimental sketch of *in vitro* differentiation of hESCs using combined TFs. (**B**) Western blot validation of protein expression of PAX6 and TFAP2A in iNPCs. (**C**) FACS analysis of induced PAX6+ and TFAP2A+ cells derived from hESCs. (**D**) GSEA cell type signature enrichments for the observed expression change following PAX6 overexpression alone. (**E**) GSEA cell type signature enrichments for the observed expression change following TFAP2A overexpression alone. (**F**) MDS (multi-dimensional scaling) visualization of different treatment groups in comparison to the target amacrine state.

**Table S1 Public datasets used in this study**.

**Table S2 Benchmark of causal discovery using simulated data with different numbers of variables and intervention fractions**.

**Table S3 Speed comparison of causal discovery in large scales**.

**Table S4 Accuracy comparison of causal discovery in large scales**.

**Table S5 Benchmark of causal discovery with Perturb-seq data**.

**Table S6 CASCADE hyperparameter robustness of causal discovery with Perturb-seq data**.

**Table S7 Benchmark of causal discovery using different scaffold graphs**.

**Table S8 Benchmark of counterfactual perturbation prediction with Perturb-seq data**.

**Table S9 CASCADE ablation test of counterfactual perturbation prediction with Perturb-seq data**.

**Table S10 CASCADE hyperparameter robustness of counterfactual perturbation prediction with Perturb-seq data**.

**Table S11 Benchmark of intervention design with held-out perturbations**.

**Table S12 Cell type signature genes used in GSEA analysis of the amacrine differentiation experiment**.

## Methods

### The generative model of CASCADE

CASCADE is a causality-aware Bayesian generative model building upon previous works of differentiable causal discovery methods^9,10,12,61^, with dedicated designs for incorporating prior regulatory knowledge, modeling genetic perturbation effects and improving computational scalability. We denote perturbational/interventional data as 𝒟 = {(**x**^(*i*)^, **r**^(*i*)^, **s**^(*i*)^)|*i* = 1, 2, …, *N*}, where **x**^(*i*)^ ∈ ℝ^*v*^, **r**^(*i*)^ ∈ {0, 1}^*v*^, **s**^(*i*)^∈ ℝ^*s*^ are the gene expression profile, interventional regime, and covariates (e.g., batch and sequencing depth) of the *i*^th^ cell. *N* is the total number of cells. *S* is the number of covariates. We denote *V* as the set of genes in the dataset. *V* = |*V*| is the total number of genes. The interventional regime 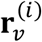 is encoded as 1 if the gene *v* ∈ *V* is intervened in the *i*^th^ cell and 0 otherwise. For convenience, the cell superscript (*i*) will be dropped when referring to any arbitrary cell below.

We model the observed data as generated by a causal regulatory graph, denoted as 𝒢 = (*V*, ℰ), ℰ ⊆ *V*×*V*, where each edge (*v*_*i*_, *v*_*j*_) ∈ ℰ represents a causal regulatory relation from gene *v*_*i*_ to gene *v*_*j*_. The adjacency matrix of 𝒢 is denoted as **A** ∈ {0,1}^*V*×*V*^ where **A**_*ij*_ = 1 corresponds to (*v*_*i*_, *v*_*j*_) ∈ ℰ. In practice, **A** is modeled as a sparse continuous logit matrix, from which binary samples are obtained using the straight-through Gumbel estimator^62^. With slight abuse of notations, we denote the set of probabilistic structural equations associated with 𝒢 as 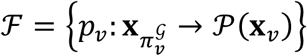 where 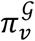 denotes the set of parent genes of gene *v*. The specific causal graph 𝒢 and structural equations ℱ are to be learned from the interventional data 𝒟.

The search space of all potential DAGs is super-exponential to the number of genes, which would be too large even for mildly large systems. To improve scalability to larger regulatory networks, we incorporate prior regulatory information as a scaffold graph 𝒢_scaffold_ = (*V*, ℰ_scaffold_) and limit the search space of causal graph 𝒢 to subgraphs of 𝒢_scaffold_.

The following unnormalized prior densities 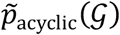 and 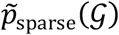 are used to introduce prior assumptions that the causal graph 𝒢 is acyclic (Eq. 1) and sparse (Eq. 2):

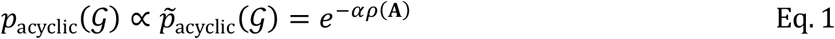

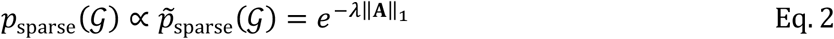

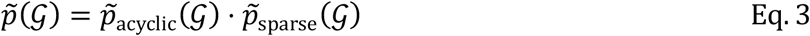

where *ρ*(**A**) is the spectral radius of the adjacency matrix **A**, serving as an approximate cycle penalty^12^, and ‖**A**‖_1_ is the *L*1 norm, serving as a sparsity penalty. *α* and *λ* are penalty coefficients that are iteratively updated during training (see below). Meanwhile, no prior assumption is imposed on the structural equations, with 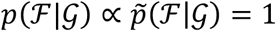.

It should be noted that gene regulation is a complex system involving extra molecular layers such as post-transcriptional and post-translational modifications, and that experimental measurements may involve uncharacterized confounding factors. As such, the above causal model defined on gene expression alone cannot capture all influencing factors. To accommodate variation unexplained by 𝒢, an intervention latent variable **z** ∈ ℝ^*Z*^ is also incorporated to accommodate residual intervention effect. We assume standard normal priors for latent **z**:

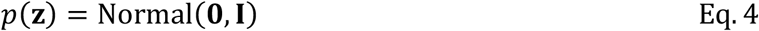

The joint distribution over data and model thus factorizes as:

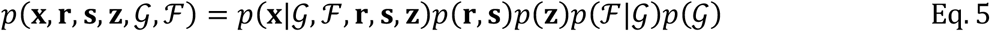

The data likelihood function can further be factorized according to the Markov properties of 𝒢:

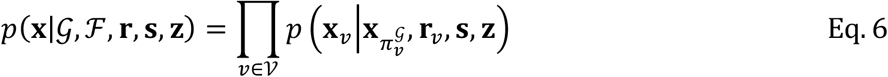

The factorized components, i.e., structural equations in ℱ, are fitted with MLPs. Two types of likelihood functions are supported, i.e., negative binomial for scRNA-seq data, and normal for simulated data.

In the case of negative binomial likelihood, the structural equations are computed as:

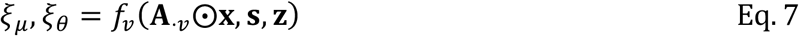

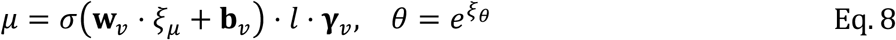

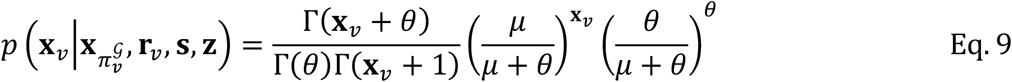

where **A**_⋅*v*_ is the *v*^th^ column of the adjacency matrix **A**, and ⨀ is the Hadamard product. *f*_*v*_ is a gene-wise structural equation in the form of an MLP. The MLP output *ξ*_*μ*_ ∈ ℝ controls the mean of negative binomial and is interpreted as the logit-scale fraction of gene *v*’s maximal count portion **γ**_*v*_ (0 < **γ**_*v*_ < 1) allowed within the cell-wise total count *l*. This formulation is chosen to roughly normalize the MLP output scales for genes with different expression levels, thus facilitating optimization. **γ**_*v*_ is determined before model fitting by raising gene *v*’s maximal count portion across all training cells to the power of −0.75, giving the MLP prediction enough head room to exceed the maximal values seen during training. *σ* is the sigmoid function converting logit to fraction. **w**_*v*_ and **b**_*v*_ are linear interventional coefficients applied on top of *ξ*_*μ*_, defined based on whether gene *v* is intervened in the cell:

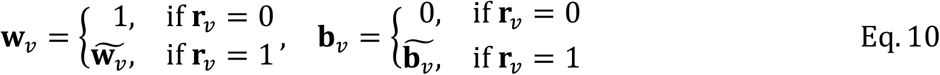

where 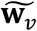 and 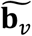 are the actual parameters to be optimized.

In the case of normal likelihood, the structural equations are computed as:

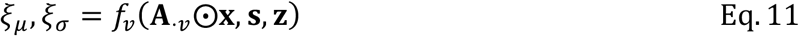

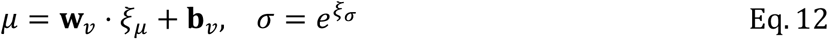

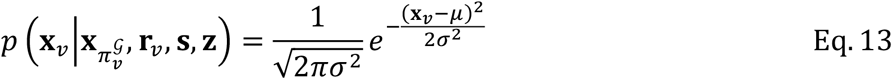

Notations are analogous to the negative binomial case.

### Posterior regulatory inference of CASCADE

We conduct posterior inference in the generative model described above to learn the causal regulatory graph 𝒢 and structural equations ℱ along with their uncertainties, which is unavoidable and intrinsic to the limited data regime found in real-word biological data. According to the Bayes rule, the posterior of 𝒢 and ℱ can be expressed as:

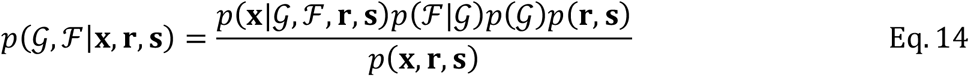

The posterior form as in Eq. 14 is intractable given that (1) *p*(**x**|𝒢, ℱ, **r, s**) implies an integral over unobserved latent **z** (Eq. 6); (2) the data marginal distributions *p*(**r, s**) and *p*(**x, r, s**) imply integrals over all possible graphs and equations; (3) the prior distributions *p*(𝒢) and *p*(ℱ|𝒢) can only be expressed in unnormalized forms (Eq. 1–Eq. 3); so approximal inference is required. Considering that edges in the graph 𝒢 must exhibit a complex dependency structure, i.e., inclusion of certain edges affects the probability of including other edges, modeling the graph posterior with parameterized distributions involving super-exponential numbers of interaction terms is unrealistic. Instead, we employ Stein variational gradient descent (SVGD)^61,63^ to obtain just posterior samples from *p*(𝒢, ℱ|**x, r, s**), which offers approximate estimates of graph uncertainty while also retaining backward compatibility with maximum *a posteriori* (MAP) estimation when drawing a single posterior sample.

SVGD is a technique that iteratively optimizes a group of *n* “particles” of causal graph and structural equation pairs {(𝒢_1_, ℱ_1_), (𝒢_2_, ℱ_2_), …, (𝒢_*n*_, ℱ_*n*_)} according to gradient of the posterior distribution and kernel-based repulsion among particles. The update gradients of the particles at each step can be expressed as:

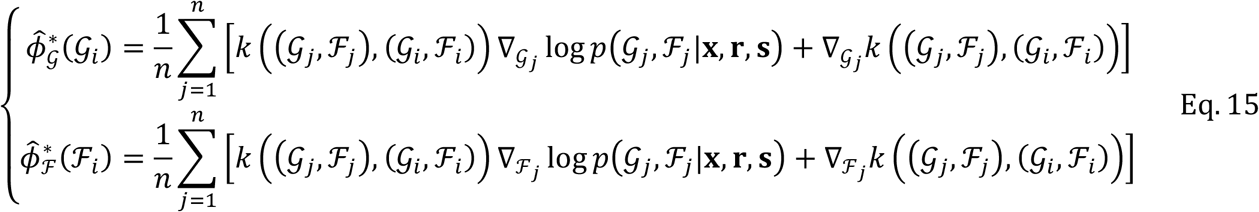

where *k*((𝒢, ℱ), (𝒢′, ℱ′)) is the kernel function. Given that the structural equations in ℱ are implemented as neural networks, evaluating their similarity could be computationally expensive. Thus, we choose to make a simplifying approximation and keep only the similarity between causal graphs, defined by the following radial basis function (RBF) kernel:

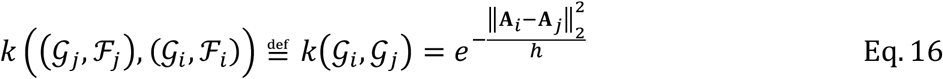

where *h* is an adaptive kernel width defined as the median of all pairwise distances among the particles. Similarly, we ignore the gradient crosstalk among particles in ℱ, and use independent updates for ℱ instead:

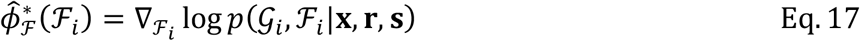

The gradient of the posterior distribution regarding 𝒢 and ℱ can be written as:

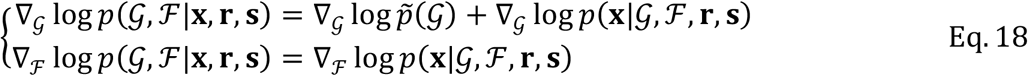

The marginal data distributions and the prior normalizing factors can be discarded due to being constants regarding 𝒢 and ℱ.

The data likelihood log *p*(**x**|𝒢, ℱ, **r, s**) is approximated using variational inference (VI)^64^ with the following evidence lower bound:

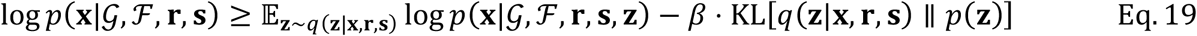

where *q* is the variational posterior of **z**, and the KL weight *β* controls how informative latent **z** can get. Since **z** is designed to accommodate residual intervention effect, we choose to simplify its variational posterior to depend only on the intervention regime:

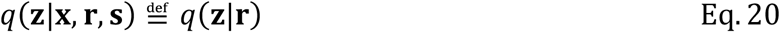

Specifically, to project intervened genes to the latent **z**, a set of biological function embeddings **u**_1_, **u**_2_, …, **u**_*V*_ ∈ ℝ^*U*^ for all genes are first obtained by performing latent semantic indexing (LSI) on a gene-by-GO annotation matrix. The function embeddings of intervened genes in a cell are merged through an attention-based pooling operation:

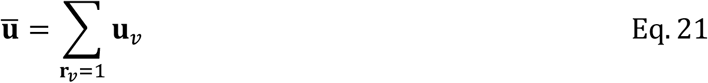

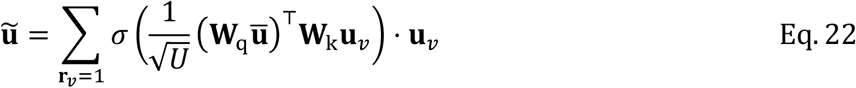

where **W**_q_, **W**_k_ ∈ ℝ^*U*×*U*^ are attention query and key projections, and *σ* is the sigmoid function. *q*(**z**|**r**) is then computed as:

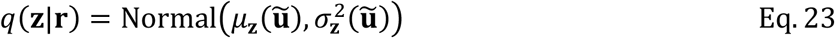

where *μ*_**z**_ and 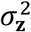 are linear transformations producing the mean and variance of the variational posterior.

The complete posterior inference procedure of CASCADE involves an iterative optimization procedure analogous to the augmented Lagrangian method^9,10,61,65^. In essence, each realization of the graph penalty coefficients (*α, λ*) (see Eq. 1, Eq. 2) defines an SVGD-VI subproblem to be optimized. We start with no penalty *α* = 0, *λ* = 0, and every time the associated subproblem converges, these coefficients are updated based on the ratios of graph gradient norms to impose increasingly strict cycle penalty, gradually pruning 𝒢 into a directed acyclic graph:

1. Randomly initialize posterior particles 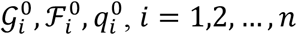, and initialize *α* = 0, *λ* = 0.
2. Iteratively update 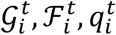 according to the following rules:

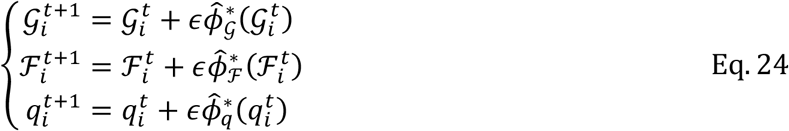

where *ϵ* is the learning rate, and

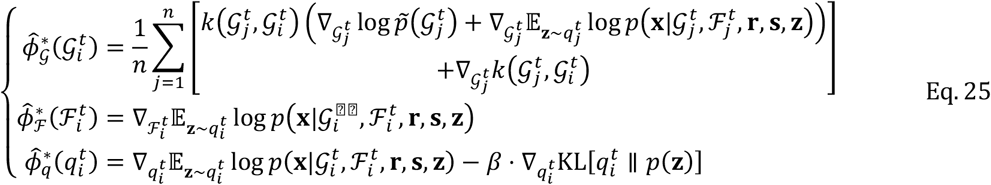
3. When the evidence lower bound converges, update the graph penalty coefficients *α* and *λ* according to the following rules:

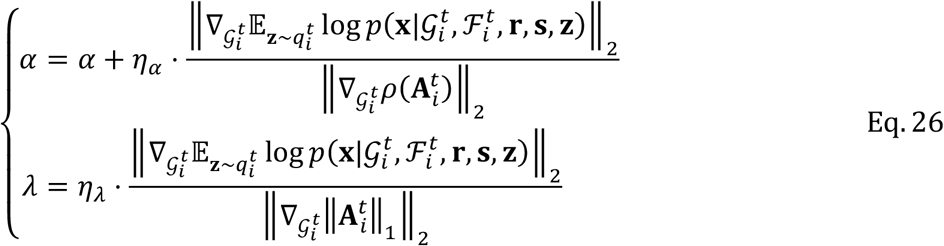

where *η*_*α*_ and *η*_*λ*_ are user specified hyperparameters controlling penalty rate.
4. Repeat step (2) until 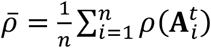 falls below a tolerance threshold.
5. Return the final posterior particles 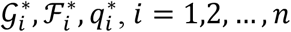.

The update rules for *α* and *λ* in Eq. 26 are formulated as such to accommodate different data that may have drastically different signal-to-noise ratios. The numerator 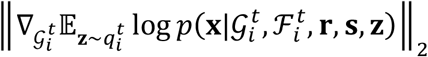 is the graph gradient norm from data likelihood, which encourages the model to preserve regulatory edges that well explain gene-gene dependency. The denominators 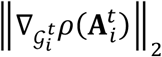 and 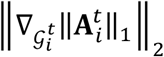 are graph gradient norms from penalties, which drive the model to break cycles and prune uninformative edges. Model training essentially strikes a balance between edge preservation and pruning. Data with higher signal-to-noise ratios produce larger likelihood gradients, which require higher penalty coefficients for effective pruning, and vice versa. It would be challenging for the user to estimate reasonable data-dependent *α* and *λ* values. So instead, they are adaptively set according to the ratio between likelihood and penalty gradient norms. We provide reasonable default values for the rate hyperparameters *η*_*α*_ and *η*_*λ*_ that are stable across different data.

While model training effectively prunes most cycles in the resulting graphs, a small number of cycles may persist due to numerical limitations. To ensure strict acyclicity required by downstream inferences, the resulting graphs 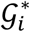 are further pruned through iteratively finding and breaking cycles by removing edges with minimal logit scores.

### Counterfactual deduction and tuning

To deduce the effect of an observed cell (**x, r, s**) under a counterfactual intervention 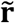 (and potentially also counterfactual covariate 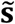) given the learned model (𝒢, ℱ, *q*), we first infer the latent variable 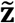 using the learned variational posterior 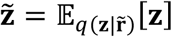. Given that all DAGs can be topologically sorted, we partition genes in *V* into topological generations *V*_1_, *V*_2_, …, *V*_*L*_ ⊆ *V* as determined by 𝒢, which satisfies 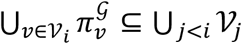, i.e., all predecessor genes of each generation are contained within previous generations. Then, the expectation of counterfactual gene expression is computed generation-by-generation:

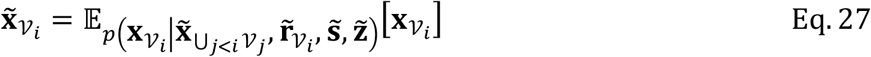

At the first generation, the predecessor set is empty ⋃_*j*<1_ *V*_*j*_ = ∅. Starting from the second generation, the predecessor genes 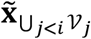 use the previously inferred counterfactual values rather than the observed values 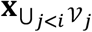. This allows intervention effect to propagate throughout the inferred causal graph 𝒢. The above computation is performed independently for each posterior particle 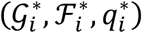, and their average counterfactual transcriptome is taken as the final prediction.

Notably, the CASCADE causal discovery step described in the previous sections is trained in a self-reconstruction setup without graph propagation, which may not seamlessly generalize to the above counterfactual setup. To further improve the model’s counterfactual accuracy, an additional fine-tuning step was devised involving random pairs of cells (**x**^(*i*)^, **r**^(*i*)^, **s**^(*i*)^) and (**x**^(*j*)^, **r**^(*j*)^, **s**^(*j*)^) in the training data. The fine-tuning objective is otherwise the same as in causal discovery, except that the self-reconstruction data likelihood is replaced by the counterfactual likelihood of cell *i* under the intervention of cell *j*, computed using the generation-by-generation graph propagation procedure implemented in an end-to-end differentiable manner. During fine-tuning, the causal graph 𝒢 is fixed, and only the structural equations in ℱ and the latent posterior *q* are updated.

### Explanation of counterfactual predictions

To dissect the contribution of different model components and upstream regulators for the counterfactual prediction of each gene *v* ∈ *V*, we estimated their contributions by independently replacing observed values with counterfactual values:

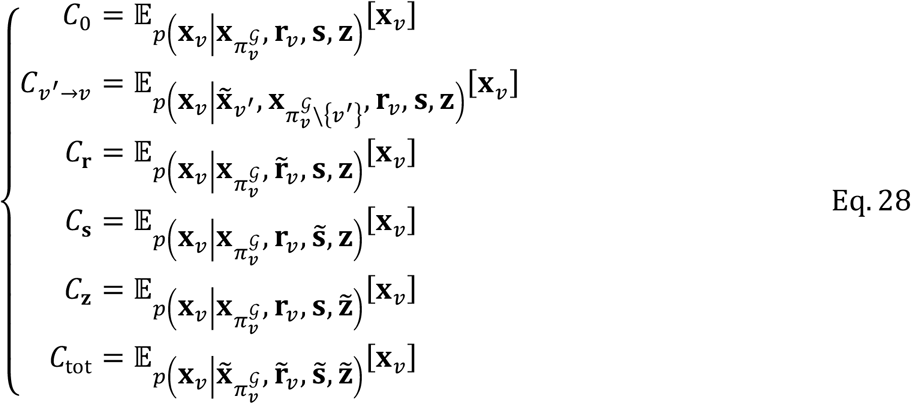

Their contribution ratios are estimated as:

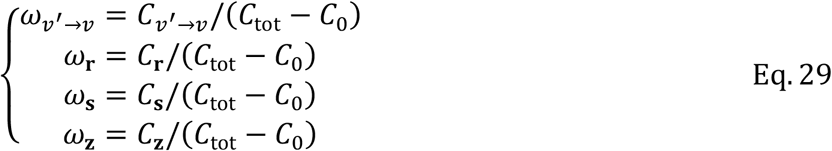

Notably, due to the non-linearity of neural networks, individual contributions are not necessarily additive, and the estimated ratios should be taken as approximations. We require total effect |*C*_tot_ − *C*_0_| to be larger than a cutoff (by default 0.1) for the above contribution analysis, which would otherwise be unreliable. The contributions are averaged across all cells with the same perturbation as well as all SVGD particles.

To explain the counterfactual prediction for a given set of response genes, we first filtered the graph 𝒢* by removing edges with *ω*_*v*′→ *v*_below a contribution cutoff (by default 0.05), producing a high contribution graph 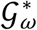. Upstream regulators of these response genes were traced via reversed traversal on 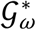 to produce a subgraph 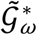, which was then visualized using Cytoscape (v3.10.3)^66^.

### Targeted intervention design

Targeted intervention design tries to find the optimal intervention 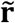, among all possible multi-gene combinatorial interventions up to a user-specified order *K*, that could produce counterfactual transcriptome 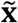 closest to a desired target transcriptome 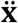. We again leverage the model’s ability to infer 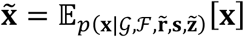 in an end-to-end differentiable manner, and minimize the following mean square error (MSE) through stochastic gradient descent:

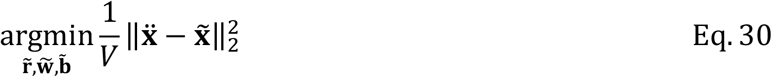

where 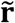 is treated as learnable rather than a fixed observed value. In practice, a logit vector 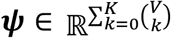 is introduced to characterize the probability of each multi-gene combination up to order *K*, and the intervention design 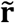 is sampled in a differentiable manner using the straight-through Gumbel estimator^62^. We also allow the intervention scales 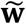 and biases 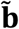 (see Eq. 10) to be optimized during the design procedure. All model parameters other than 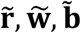 are held constant.

### Implementation details

The CASCADE model was implemented in PyTorch. Each structural equation in ℱ is implemented as an MLP containing a single 16-dim hidden layer, leaky-ReLU activation and 20% dropout ratio. The fully connected layers across all MLPs are combined into a single batched matrix multiplication to ensure computational efficiency on GPUs. While Eq. 7 and Eq. 11 suggest using the entire column vector **A**_⋅*v*_ to mask non-parent genes in **x**, both **A**_⋅ *v*_ and **x** are first condensed and zero-padded to the maximal in-degree of the scaffold graph (usually much smaller than *V*), so as to reduce the input dimension of the neural networks and maximize computational efficiency. Default values of other model hyperparameters include the number of SVGD particles *n* = 4, latent **z** dimension *Z* = 16, KL weight *β* = 0.1, graph penalty rates *η*_*α*_ = 0.5, *η*_*λ*_ = 0.1.

Across causal discovery, counterfactual tuning and intervention design, we use 90% of the data for training and 10% for validation. Batch size is set as 128. Learning rate *ϵ* is 0.005 for causal discovery, 0.0005 for counterfactual tuning and 0.05 for intervention design. The validation loss is checked every 300 training steps. Early stopping is triggered when validation loss no longer decreases for a consecutive 3 loss checks. The tolerance for cycle penalty 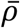 is 0.0001. All CASCADE models in the manuscript use the above default setup unless otherwise stated.

### Data simulation for causal discovery benchmarks

Given a gene number *V*, the underlying true DAG 𝒢 was first generated by randomly sampling 10% of the entries in the upper triangular part of the adjacency matrix. This ensures that the resulting graph is directed and acyclic. We then randomly assign weights of 1 or −1 to the sampled edges, resulting in an signed adjacency matrix **A** ∈ {−1,0,1}^*v*×*v*^. Following the topological generations (*V*_1_, *V*_1_, …) of graph 𝒢, we iteratively simulate the values of all genes based on the following structural equation:

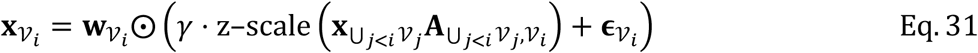

where **ϵ** ∈ ℝ^*V*^ is the exogenous noise sampled from the standard normal distribution Normal(**0, I**_*V*_), *γ* controls the signal-to-noise ratio, and 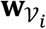 are intervention coefficients dictating whether and how each gene is intervened in the cell. ⋃_*j*<*i*_ *V*_*j*_ represents all topological generations prior to the current one being simulated. Z-scaling is included to maintain constant marginal gene variances and prevent them from being correlated with the causal order, which certain causal discovery methods might otherwise exploit^16^.

In all benchmarks, we use *γ* = 5, and “knockout” intervention coefficients of **w**_*i*_ ∈ {0,1}. Intervened genes are uniformly sampled from all genes. We always simulate a total of 50,000 cells, and the numbers of cells in each interventional regime (including the non-interventional case) are uniformly distributed.

### Benchmarking with simulated data

PC, GES, and GIES were executed using the R package “pcalg” (v2.7-12). ICP, NOTEARS, DAGMA, NO-BEARS, DCDI and DCD-FG were executed using Python packages “causalicp” (v0.1.1)^67^, “notears” (v3.0)^9^, “dagma” (v1.1.1)^65^, “BNGPU” (commit d4c6f48)^12^, “dcdi” (commit 594d328)^10^ and “dcdfg” (commit 7ad08e8)^11^, respectively. All methods were run using their default hyperparameters, on Linux servers equipped with 28 CPU cores (2 Intel Xeon Gold 6132 chips), 192GB RAM and NVIDIA V100 GPUs (32GB VRAM). Only a single GPU was used for applicable methods. CASCADE was run using the normal likelihood and a single SVGD particle (*n* = 1).

Structural Hamming distance (SHD) was computed as the L1 distance between inferred adjacency matrix and the true adjacency matrix, i.e., edge reversals are counted as 2:

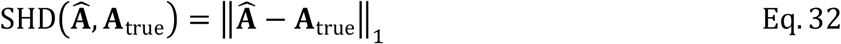

Average precision, AUROC, precision, and recall were computed only among edges that exist in the scaffold graph, using the Python package “scikit-learn” (v1.5.1).

### Construction of the scaffold graph

Given that CASCADE is more sensitive to false negative than false positive edges in the scaffold graph (Fig. 2D, Fig. S3), we sought to build a scaffold graph for real biological data by integrating 4 heterogeneous forms of prior regulatory knowledge in a context agnostic manner in order to cover as many potential edges as possible and minimize false negatives. Specifically, we combined gene-gene relations in KEGG pathways^19^, protein-protein interactions in the BioGRID database^22^, gene expression correlation in the GTEx data^23^, and TF-target gene pairs bridging ENCODE TF ChIP-seq bindings^20^ with cis regulatory elements (CRE) annotations in the GeneHancer database^21^.

Incorporating KEGG and BioGRID edges was relatively straightforward. For GTEx data, we computed the gene-gene correlation matrix across 19,616 samples, and binarized it into a graph by taking gene pairs that are mutually within the top 500 correlated genes. For TF-target gene pairs, we took the irreproducible discovery rate (IDR) thresholded peaks from all ENCODE human TF ChIP-seq data and intersected them with target gene-annotated enhancers in GeneHancer and gene promoters (–2,000 to +500 bp from transcription start sites), producing a TF-by-target intersection count matrix. To account for different enhancer lengths and ChIP-seq sensitivities, the intersection counts were normalized by expected values from row and column marginals. Finally, we compared the normalized counts with a null distribution obtained from randomly permuted matrices and retained TF-target pairs with a permissive empirical *P*-values < 0.15.

The number of edges in each prior knowledge type were: 83,825 for KEGG (directed), 878,248 for BioGRID (undirected), 1,838,142 for GTEx (undirected) and 1,629,136 for TF-target gene (directed). The final scaffold graph was constructed by taking the union across all 4 edge types.

### Perturb-seq data preprocessing

Cells with perturbation of genes absent from the readout profile were first discarded. The raw expression counts were log-normalized for causal discovery and perturbation prediction methods not supporting count input. Highly variable genes were identified and ranked using the “sc.pp.highly_variable_genes” function in “scanpy” (v1.10.2)^68^. Cells with perturbation label but no perturbation effect were removed using “pt.tl.Mixscape” in “pertpy” (v0.8.0)^69,70^. Finally, perturbations with less than 10 cells were also removed. The global effect size of each perturbation was quantified with E-distance using the “edist_to_control” function in “scperturb” (v0.1.0)^71^.

To allow models to account for technical confounding, we set up covariates for each dataset as follows. Given that several Perturb-seq datasets involve a large number of batches^18^, we applied singular value decomposition (SVD) on the batch-averaged expression matrices to obtain low-dimensional batch embeddings. The SVD dimensions were chosen based on the elbow point of explained variance ratios. The resulting batch embedding, log-transformed total count per cell and a binary “is perturbed” flag was concatenated and used as the covariate.

### Benchmarking unseen perturbation prediction and intervention design with independent Perturb-seq data

The Perturb-seq datasets were divided into training and test data as described in GEARS^26^. All perturbed genes were first randomly partitioned into *V*_seen_ and *V*_unseen_ subsets at a 9:1 ratio. Single-gene perturbations in *V*_seen_ were assigned to the training set, and those in *V*_unseen_ to the test set. Multi-gene perturbations consisting entirely of *V*_seen_ genes were randomly split into training and test sets at a 3:1 ratio, while those containing any gene in *V*_unseen_ were all assigned to the test set.

To increase signal-to-noise ratio, nearest neighbor-based data imputation was conducted. Specifically, *k*-nearest neighbor search was conducted using PCA embeddings among cells of the same perturbation labels. The gene expression profile of each cell was then imputed by aggregating the expression in its neighbors (*k* = 20 by default).

The baseline methods “Mean”, “Additive” and “Linear” were implemented as described in an independent benchmark study^31^. For all three baselines, training cells receiving the same perturbations were first aggregated by taking the average of their log-normalized transcriptomes to produce **x**^(*v*)^(*v* ∈ *V*_train_ ⊆ *V* denotes the perturbed gene). The “Mean” baseline then predicts the mean across all training perturbations 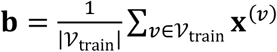 regardless of what perturbation is being predicted. The “Additive” baseline first computes the perturbational change *δ***x**^(*v*)^= **x**^(*v*)^− **x**^∅^(**x**^∅^ is the unperturbed state) for each training perturbation. For each perturbation *r* ⊆ *V* to be predicted (may involve multiple genes), the prediction is then 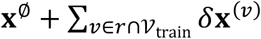, i.e., For *v* ∈ *r* not in the training set, the corresponding change *δ***x**^(*v*)^ is simply dropped. The “Linear” baseline first conducts a *k*-dimensional principal component analysis (PCA) over the training perturbation change matrix 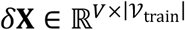 to produce low-rank embeddings for each readout gene **G** ∈ ℝ^*V*×*k*^. The rows in **G** corresponding to perturbed genes were extracted as a perturbation embedding 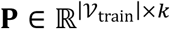. A bilinear linear regression for *δ***X** was solved in closed form:

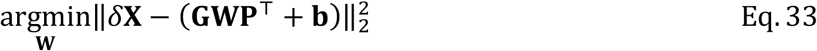

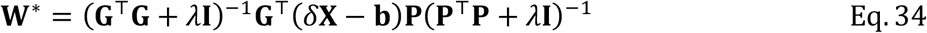

where *λ* is the ridge penalty. For perturbation *r* ⊆ *V* to be predicted (may involve multiple genes), the perturbation changes for each *v* ∈ *r* were predicted by taking the corresponding gene embedding in **G** as the perturbation embedding **p**^(*v*)^, and computing 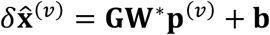. The final prediction is then 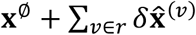. The default of *k* = 10 and *λ* = 0.1 were used as described in the benchmark study^31^.

CPA, Biolord, GEARS, scGPT, scFoundation, STATE were executed using Python packages “cpatools” (v0.8.7)^24^, “biolord” (v0.0.3)^25^, “cell-gears” (v0.1.1)^26^, “scgpt” (commit 706526a)^28^, “scFoundation” (commit d09626a)^29^ and “state” (commit 0221d05)^30^ respectively. We used the same GO LSI embeddings used in CASCADE for CPA and Biolord. For scGPT and scFoundation, we initialized the models using the pretrained weight publicly released by the authors and further fine-tuned them on the benchmark datasets. All methods were run using their default hyperparameters, on Linux servers equipped with 64 CPU cores (2 Intel Xeon Platinum 8358 chips), 1TB RAM and NVIDIA A100 GPUs (80GB VRAM). Only a single GPU was used for applicable methods. CASCADE was run using the negative binomial likelihood.

To compute “responsiveness accuracy”, we first assessed the perturbation effect for each inferred edge. For each edge in the inferred graph (*v*_*i*_, *v*_*j*_) ∈ ℰ where both *v*_*i*_ and *v*_*j*_ are perturbed in the test data, we compared expression change of *v*_*j*_ upon *v*_*i*_ perturbation (forward direction) and expression change of *v*_*i*_ upon *v*_*j*_ perturbation (reverse direction), using the Kolmogorov-Smirnov test. Denote *n*_fwd_ as the number of edges where forward *P*-value < 0.1 and reverse *P*-value > 0.1, and *n*_rev_ as the number of edges where reverse *P*-value < 0.1 and forward *P*-value > 0.1. Responsiveness accuracy is then defined as:

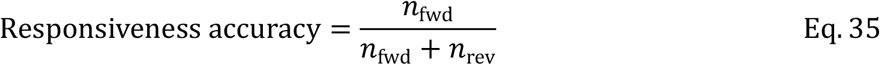

For each perturbation, delta correlation and normalized MSE were computed using average expression profiles in predicted counterfactual cells 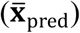, true perturbed cells 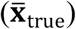 and control cells 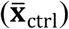, defined as follows:

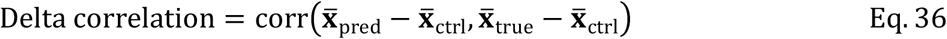

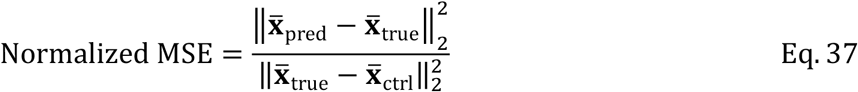

The scores reported in Fig. 3A, D, Fig. S9, Fig. S10 were average delta correlation and normalized MSE across all test perturbations in each dataset/category that have E-distance > 2.

Aside from CASCADE, we also adapted the above perturbation response prediction methods to perform intervention design, or “reverse perturbation prediction”, using the ensemble voting approach as proposed in scGPT^28^. Briefly, for each candidate perturbation *r*_*i*_ ⊆ *V, i* = 1,2, …, *R*, we predict the perturbed transcriptome for 30 randomly sampled control cells to form a reference panel (of size 30 ⋅ *R*). Each ground truth perturbed cells (assuming true perturbation is 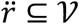) is then used to query the reference panel for nearest neighbors and count their perturbations as votes. Vote counts *n*_1_, *n*_2_, …, *n*_*R*_ are taken as design scores for the perturbation 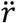. Note that the vote count vector ***n*** = [*n*_1_, *n*_2_, …, *n*_*R*_] is analogous to the design score vector ***ψ*** in CASCADE and will be used interchangeably below.

To quantify design accuracy, we define hit rate curve (HRC) as follows. For each ground truth perturbation 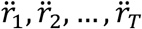, we obtain the quantile 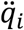of 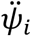 among ***ψ***_***i***_ sorted from largest to smallest, so 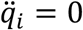 indicates that the design score of the true perturbation 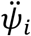 is the largest among all candidate perturbations for ground truth 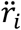. We then sort 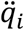 ‘s of all *T* ground truth perturbations from smallest to largest and assign hit rates 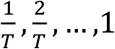, which indicates that if we threshold design quantile at 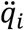, we would expect to hit 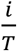 of ground truth perturbations. The HRC is thus the curve with design quantiles on the x-axis and hit rate on the y-axis, analogous to an ROC curve (also see Fig. S11A for an illustration). The area under hit rate curve (AUHRC) was computed using the “auc” function in “scikit-learn”. For each test split of each dataset, we chose the *T* = 5 perturbations with largest E-distance as design targets. Larger AUHRC indicates more accurate intervention designs. Fig. 4A was produced with all test splits of all Perturb-seq datasets combined.

### Intervention design of erythroid differentiation

The “Norman-2019” K562 CRISPRa Perturb-seq dataset^17^ was used as training data. Five single-cell datasets containing erythroid cells were used as the design target, i.e., “Xu-2022”^32^, “Huang-2020”^33^, “Xie-2021”^34^, “Multiome-2021”, and “CITE-2021”^35^. For the multi-omics datasets, only the RNA modality was used. For datasets contain additional lineages, only the erythroid cell types (“Proerythroblast”, “Erythroblast”, “Normoblast”, “Reticulocyte”) were used. Genes were selected for model training if they (1) were among the top 5,000 highly variable genes in “Norman-2019”, and (2) exhibit differential expression between K562 and erythrocytes, or (3) were perturbed in the Perturb-seq dataset. A total of 1,147 genes were selected. CASCADE was trained on the “Norman-2019” dataset using the heterogeneous scaffold graph. The covariates were set as described in “Perturb-seq data preprocessing”.

For erythroid targeted intervention design, we selected genes that were (1) perturbed in “Norman-2019”, and (2) exhibit higher expression in erythroid cells as the candidate set, resulting in 32 candidate genes. The covariates for target erythroid cells were imputed by using the covariates of the most similar “Norman-2019” cells in the space of stable housekeeping genes. To allow the model to focus on the modulation of erythroid genes, we compiled a list of 70 known erythroid markers. The design MSE loss in Eq. 30 was computed in a weighted manner such that the total weight of erythroid markers equals the total weight of remaining genes.

### Intervention design of targeted hESC differentiation

The H1 TF atlas dataset^40^ was used as the primary training data. To enrich for effective perturbations, only perturbed cells in the author annotated clusters 6, 7, 8 exhibiting differentiation tendencies were used, along with control cells in all clusters. The human fetal gene expression atlas^41^ was used as the design target. To balance cell type populations, all cell types were subsampled to a maximum of 5,000 cells. Genes were selected for model training if they (1) were highly variable in the H1 dataset (following author provided criteria “min_mean=0.0125, max_mean=3, min_disp=0.5”), and (2) exhibit differential expression between H1 and at least 50% of target fetal cell types, or (3) were perturbed in the H1 TF atlas. A total of 3,777 genes were selected. For H1 cells, covariates were set as described in “Perturb-seq data preprocessing”. For target fetal cells, the covariates were imputed by using the covariates of the most similar H1 cells in the space of stable housekeeping genes. CASCADE was trained on a combined dataset containing both H1cells and a subsampled 50,000 fetal cells, with an additional binary batch covariate to distinguish H1 and fetal cells, and the heterogeneous biological scaffold graph.

Intervention designs were conducted for all 62 well defined fetal cell types (excluding 15 only named after markers). We selected genes that were (1) perturbed in the TF atlas, and (2) exhibit higher expression in the target cell type as the candidate set. To allow the model to focus on the modulation of target cell type markers, we compiled a list of known markers for each cell type (Supplementary Table TODO). The design MSE loss in Eq. 30 was computed in a weighted manner such that the total weight of target cell type markers equals the total weight of remaining genes. For each cell type, we conducted intervention designs with up to 3-gene combinations and manually selected the most parsimonious intervention that received high design score and produced low MSE from the target cell type. Cell type signature enrichment analysis was conducted using gseapy (v1.1.3)^72^ over “c8” gene sets on MSigDB^73^, with genes ranked by log fold change of CASCADE counterfactual predictions given the designed interventions.

### Cell culture and differentiation

For routine maintenance, the human embryonic stem cell (hESC) line WIBR3 was cultured in Essential-8™ medium (Gibco, A1517001) on Matrigel™ (Corning, 354277)-coated feeder-free plates. Cells were passaged every 5–6 days using Versene Solution (Gibco, 15040066). The ROCK inhibitor Y-27632 (10 μM; Selleck, S1049) was added to the culture medium for the first 24 hours following thawing or passaging. Mycoplasma contamination was routinely monitored, and all tests are negative results.

Induced neural progenitor (iNP) differentiation was performed based on a previously described protocol^40^. Briefly, hESC culture medium was gradually transitioned to neural progenitor (NP) medium in 25% increments beginning on Day 0. NP medium was composed of DMEM/F-12 (Gibco, Cat. No. 11320082), B-27 supplement (Gibco, 17504044), 20 ng/mL EGF (Novoprotein, C029), 20 ng/mL bFGF (Novoprotein, GMP-C046), and 2 mg/mL heparin (Sigma, H3149). The medium was fully replaced with 100% NP medium at day3, and cells were passaged on day 4.

To assess the effects of transcription factor (TF) overexpression (OE) during iNP differentiation, TF OE plasmids were transiently introduced into iNPCs using the Neon™ NxT Electroporation System with the following parameters: 1,050 V pulse voltage, 30 ms pulse width, and 2 pulses. Following electroporation, cells were cultured in NP medium for an additional 3 days prior to fluorescence-activated cell sorting (FACS). EGFP^+^/ECFP^+^ cells were sorted as OE populations. Sorted cells were resuspended in DPBS supplemented with 0.04% BSA and immediately processed for downstream RNA-seq library preparation.

### Construction of the overexpression plasmids

OE constructs were generated using the ClonExpress II One Step Cloning Kit (Vazyme, C115). The coding sequences of human PAX6 and TFAP2A were individually cloned into the pPBbsr2 vector, fused with either a Flag or HA epitope tag, and co-expressed with EGFP or ECFP, respectively. In both constructs, the fluorescent protein was co-translated with the target TF and separated by a P2A self-cleaving peptide. All plasmids were confirmed by Sanger sequencing prior to use.

### Bulk RNA sequencing

Total RNA from PAX6^+^/TFAP2A^+^ cells was harvested using the Quick-RNA Microprep Kit (Zymo Research, R1051). Subsequently, 100-300 ng of the total RNA was subjected to ribosomal RNA depletion using the Ribo-MagOff rRNA Depletion Kit (Vazyme, N420). The resulting rRNA-depleted RNA was utilized for RNA sequencing library preparation using the VAHTS Universal V10 RNA-seq Library Prep Kit (Vazyme, NR606) according to the manufacturer’s protocol. Sequencing was performed on the Illumina NovaSeq X Plus by Annaroad Gene Technology (Beijing, China).

### Bulk RNA-seq data analysis

Paired-end RNA-seq reads were processed with fastp (v0.23.2) for quality filtering and adapter trimming. Processed reads were aligned to the human reference genome (GENCODE v37, primary assembly) using STAR (v2.7.9a), with only uniquely mapped reads retained for downstream quantification. Gene-level counts and transcript-per-million (TPM) values were generated using featureCounts (Subread v2.0.1) based on the GENCODE v37 gene annotation. Differential expression analysis was conducted using Limma (v3.58.1), comparing OE groups with iNPC at day 7 as the negative control. GSEA analysis was conducted with gseapy (v1.1.3) using Limma-estimated log fold changes. Cell type signature gene sets were defined as in Table S12.

## Data availability

All datasets used in this study were previously published and were obtained from public data repositories, as summarized in Table S1. Source data underlying benchmark results are available in Table S2–11. Raw and processed RNA-seq data in amacrine differentiation validation are available at GEO (accession no. GSE305268).

## Code availability

The CASCADE model was implemented in the “cascade-learn” Python package, which is available at https://github.com/gao-lab/CASCADE. For reproducibility, the scripts for all benchmarks and case studies were assembled using Snakemake (v8), which is also available in the above repository.

